# AI-Enhanced RAIN Protocol: A Systematic Approach to Optimize Drug Combinations for Rectal Neoplasm Treatment

**DOI:** 10.1101/2024.05.28.596215

**Authors:** Nasrin Dashti, Ali A. Kiaei, Mahnaz Boush, Behnam Gholami-Borujeni, Alireza Nazari

## Abstract

**Background:** Rectal cancers, or rectal neoplasms, are tumors that develop from the lining of the rectum, the concluding part of the large intestine ending at the anus. These tumors often start as benign polyps and may evolve into malignancies over several years. The causes of rectal cancer are diverse, with genetic mutations being a key factor. These mutations lead to uncontrolled cell growth, resulting in tumors that can spread and damage healthy tissue. Age, genetic predisposition, diet, and hereditary conditions are among the risk factors. Treating rectal cancer is critical to prevent severe health issues and death. Untreated, it can cause intestinal blockage, metastasis, and deteriorate the patient’s quality of life. Effective treatment hinges on finding the right drug combinations to improve therapeutic outcomes. Given the intricacies of cancer biology, treatments often combine surgery, chemotherapy, and radiation, with drugs chosen to target different tumor growth mechanisms, aiming to reduce the tumor and limit side effects. The continuous advancement in cancer treatments highlights the need for ongoing research to discover new drug combinations, offering patients improved recovery prospects and a better quality of life. This background encapsulates a detailed yet succinct understanding of rectal neoplasms, their origins, the urgency of treatment, and the quest for effective drug therapies, paving the way for discussions on treatment advancements and patient care impacts.

**Method:** This study employed the RAIN protocol, comprising three steps: firstly, utilizing the GraphSAGE model to propose drug combinations for rectal neoplasm treatment Each node in the graph model is a drug or a human gene/protein that acts as potential target for the disease, and the edges are P-values between them; secondly, conducting a systematic review across various databases including Web of Science, Google Scholar, Scopus, Science Direct, PubMed, and Embase, with NLP investigation; and thirdly, employing a meta-analysis network to assess the efficacy of drugs and genes in relation to each other. All implementations was conducted using Python software.

**Result:** The study evaluated the efficacy of Oxaliplatin, Leucovorin, and Capecitabine in treating Rectal Neoplasms, confirming their effectiveness through a review of 30 studies. The p-values for individual drugs were 0.019, 0.019, and 0.016 respectively, while the combined use of all three yielded a p-value of 0.016.

**Conclusion:** Given the significance of rectal neoplasms, policymakers are urged to prioritize the healthcare needs of affected individuals. Utilizing artificial intelligence within the RAIN protocol can offer valuable insights for tailoring effective drug combinations to better address the treatment and management of rectal neoplasms patients.

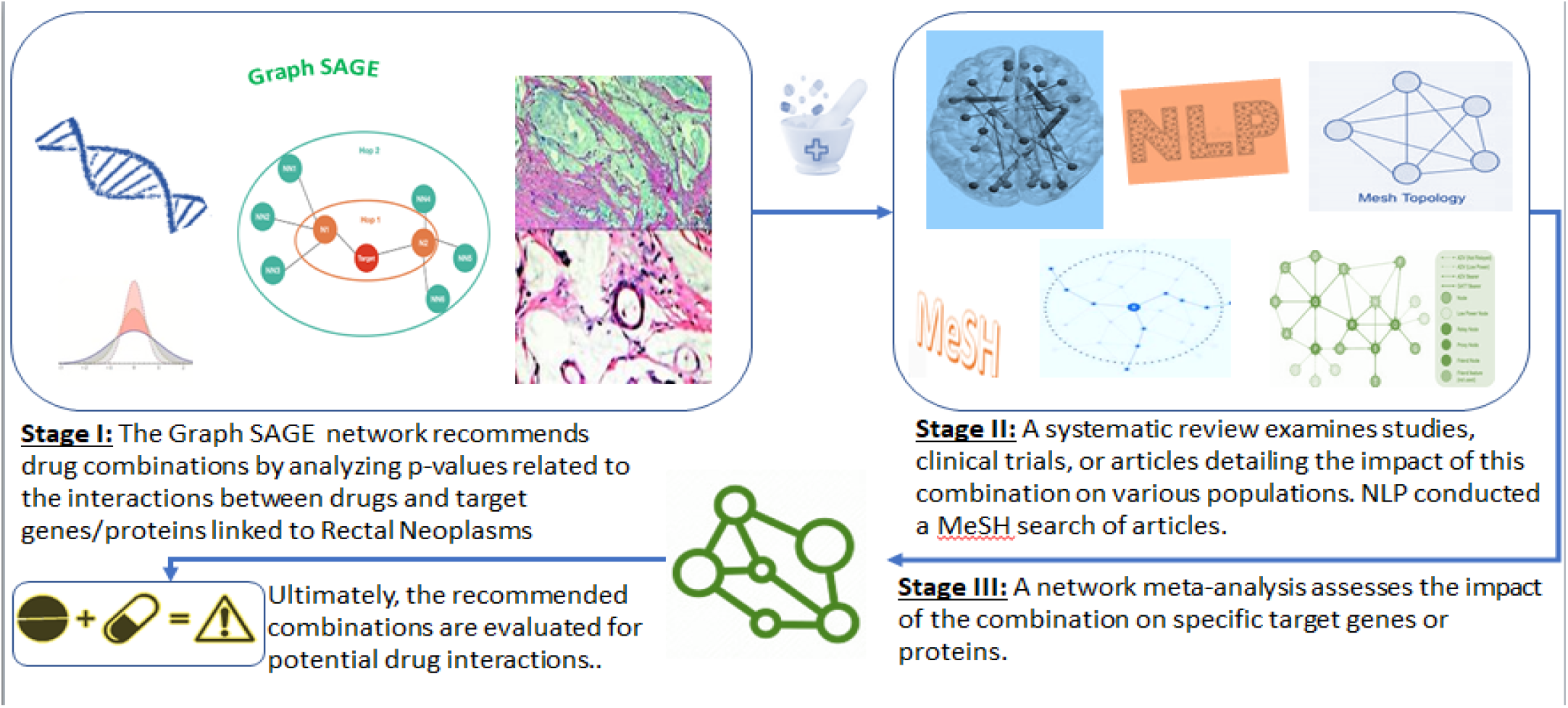

**Highlights:**

- Rectal cancers, evolving from benign polyps to malignancies, underscore the critical need for timely and effective treatment to prevent severe health complications.
- Genetic mutations, a pivotal factor in rectal cancer, trigger uncontrolled cell growth and necessitate targeted drug therapies to combat tumor spread.
- The RAIN protocol, leveraging the GraphSAGE model and systematic reviews, offers a novel approach to identify potent drug combinations for rectal neoplasm treatment.
- The study’s findings advocate for policy intervention to ensure that healthcare systems adequately support individuals battling rectal neoplasms, with AI-driven protocols enhancing patient care.

## 1. INTRODUCTION

According to Frank Gaillard, rectal neoplasms, or rectal cancer, originate in the rectum, which is the terminal portion of the large intestine. This condition begins where the final part of the colon concludes and extends to the narrow passage leading to the anus. Both rectal cancer and colon cancer collectively constitute colorectal cancer. Symptoms of rectal neoplasms may include alterations in bowel habits such as diarrhea, constipation, or frequent bowel movements, presence of dark red or bright red blood in the stool, loose stools, sensation of incomplete bowel emptying, abdominal pain, unexplained fluctuations in weight, weakness, or fatigue. (1) Effective management of rectal neoplasms plays an essential role in improving patient outcomes and overall quality of life. This body of literature provides valuable insights into various aspects of the management of rectal neoplasms. For example, Mutch & Hunt’s 2023 research emphasizes the continued importance of short-term chemotherapy (TNT) in the treatment of rectal cancer and highlights the importance of established therapies. Furthermore, Krasnick’s 2023 study sheds light on the complex nature of rectal cancer management in the context of Lynch syndrome and emphasizes the need for a multidisciplinary approach to address the clinical complexities associated with this disease. Addressing the management of intestinal neoplasms requires a comprehensive strategy that prioritizes nerve preservation, timely interventions, and personalized care, all with the goal of increasing patient well-being and survival. (2),(3) In addition, Zhou et al.’s 2022 study emphasizes the importance of sclerotherapy as a vital method for the effective treatment of rectal prolapse in pediatric patients and emphasizes the necessity of diverse approaches tailored to different patient demographics. Similarly, Alrahawy et al.’s 2022 review emphasizes the importance of careful management in rectal cancer cases, and highlights the potential of tissue analysis as a prognostic biomarker, thereby supporting the development of personalized treatment regimens. he does. In addition, a 2023 study by Zeng et al shows the clinical relevance of lymph node ratio and blood parameters in predicting recurrence-free survival among patients with high-grade rectal neuroendocrine neoplasms, emphasizing the utility of prognostic indicators in clinical decision-making. he does. Collectively, these findings emphasize the multifaceted nature of the management of rectal neoplasms, which includes elements such as early diagnosis, appropriate treatment approaches, multidisciplinary collaboration, and individualized patient care strategies. Overall, these insights underscore the critical importance of comprehensive and tailored management strategies in optimizing outcomes for individuals with rectal neoplasms(4),(5),(6) Recent research shows that drug combinations have promising benefits for the treatment of rectal neoplasms, as shown in the 2024 study by Yu Song et al. And their findings suggest that incorporating bevacizumab (BEV) into treatment regimens can increase antitumor efficacy, thereby improving disease control and managing metastatic colorectal cancer (mCRC) more effectively. BEV exhibits superior antitumor activity compared to individual therapies, especially when combined with other therapies, leading to increased disease control for mCRC patients. This integrated approach shows potential for optimizing disease management. In addition, adjunctive drugs such as cetuximab and panitumumab may further enhance treatment outcomes when combined with BEV, creating a multipronged attack against cancer cells. Lynn Eldridge’s 2024 research supports the idea that drug combinations reduce the likelihood of tumor resistance, similar to how antibiotic combinations fight resistant bacteria. In addition, the synergistic effects caused by the simultaneous use of drugs can enhance the therapeutic benefits and surpass the effectiveness of each drug alone. Early administration of combination therapy may improve treatment efficacy and disease progression, while using lower doses of drug combinations may reduce drug side effects without compromising efficacy.(7),(8)

### 1.1. Associated human genes

#### TRNAG1

Scientists have pinpointed predictive biomarkers linked to colorectal cancer (CRC) as potential treatment focal points. Pouria Samadi and collaborators identified differentially expressed genes (DEGs) through meticulous data analysis, uncovering 37 genes conserved across all CRC stages, pivotal in CRC development. Additionally, they highlighted differentially expressed micro-RNAs (miRNAs) and long non-coding RNAs (lncRNAs) associated with these DEGs. Notably, the lncRNA LINC00974 emerged as a significant and novel biomarker with potential for clinical prognosis and tumor growth/metastasis diagnosis in CRC. Future research may explore LINC00974 as a therapeutic target. Thus, comprehending molecular mechanisms involving genes like LINC00974 promises valuable insights into CRC pathogenesis across all stages.(9),(10),(11)

#### TRG-TCC1-1

In a study by Xuan-zhang Huang et al., which examined the relationship between tumor regression grade (TRG), pathologic complete response (pCR), and long-term survival in rectal cancer after neoadjuvant therapy, it was found that patients responded Pathological completeness (pCR) after neoadjuvant therapy showed significantly superior overall survival (OS) and disease-free survival (DFS) compared to those without pCR. The 10-year OS rate was significantly higher in the pCR group at 80.5% compared to 48.3% in the non-pCR (npCR) group, with a 5-year DFS rate of 90.1% in the pCR group versus 69.8% in the npCR group. The group further observed that improved tumor regression after neoadjuvant therapy was associated with improved OS and DFS rates, suggesting that better tumor regression is associated with favorable survival outcomes. While this study does not specifically address TRG-TCC1-1, it emphasizes the importance of lncRNA LINC00974 as a potential therapeutic target and prognostic indicator for rectal cancer diagnosis and prognosis, and its relevance for further exploration. It shows as a therapeutic target.

Understanding the broader molecular mechanisms, including genes such as LINC00974, promises to provide valuable insights into the pathogenesis of rectal cancer and potential therapeutic avenues.(12),(13)

#### TRG-GCC2-4

In the studies on rectal cancer by Xuan-zhang Huang and colleagues, the term TRG-GCC2-4 was not directly linked with any specific gene or biomarker. However, we draw insights from pertinent research findings in this domain, wherein tumor regression grade (TRG) denotes the evaluation of tumor response to neoadjuvant therapy among rectal cancer patients. TRG categorizes the extent of tumor regression post-treatment, with a higher TRG indicating a more favorable response and potentially enhanced survival rates. Pathological complete response (pCR), defined as the total eradication of tumor cells following neoadjuvant therapy, correlates with improved overall survival (OS) and disease-free survival (DFS) in rectal cancer patients. Moreover, attention has been drawn to the significance of lncRNA LINC00974 as a promising therapeutic target, serving as a prognostic indicator for clinical diagnosis and tumor growth/metastasis assessment in rectal cancer. Future investigations may delve into the therapeutic potential of LINC00974. A comprehensive grasp of the molecular mechanisms involving genes like LINC00974 promises valuable insights into rectal cancer pathogenesis and potential therapeutic interventions.(12),(14),(15)

#### TRG-CCC2-1

In the investigations into rectal cancer conducted by Xuan-zhang Huang and colleagues, the term TRG-CCC2-1 did not appear to have a direct association with any specific gene or biomarker. Nevertheless, we offer insights gleaned from pertinent research in this domain concerning tumor regression grade (TRG), which evaluates the response of rectal cancer tumors to neoadjuvant therapy. This assessment classifies the extent of tumor regression post-treatment, with a higher TRG indicating a more favorable response and potentially improved survival outcomes. Moreover, achieving pathologic complete response (pCR), characterized by the complete disappearance of tumor cells following neoadjuvant therapy, has been linked to enhanced overall survival (OS) and disease-free survival (DFS) in patients with rectal cancer. Notably, emphasis has been placed on the significance of lncRNA LINC00974 as a promising therapeutic target, serving as a marker for clinical prognosis and the diagnosis of tumor growth and metastasis in rectal cancer. Further exploration may delve into the potential therapeutic role of LINC00974. A comprehensive comprehension of the molecular mechanisms involving genes like LINC00974 promises valuable insights into the pathogenesis of rectal cancer and potential therapeutic approaches (15),(16)

#### LARS

Low anterior resection syndrome (LARS) manifests as a cluster of symptoms that may arise post-surgery, particularly after the removal of the lower part of the colon, notably the rectum. LARS encompasses symptoms involving stool-related issues following procedures like rectal sphincter removal, such as incontinence characterized by the inability to control bowel movements, leading to involuntary stool leakage, as well as stool urgency, marked by sudden and intense urges to defecate. Dysfunction of the rectum further compounds defecation problems. These symptoms significantly impact the patient’s quality of life. Historically, abdominoperineal resection (removal of the rectum and anus) was the standard treatment for lower rectal cancers. However, advancements in surgical techniques, equipment, and neoadjuvant therapy have made sphincter-preserving methods the preferred approach. The objective of sphincter-preserving techniques is to achieve a negative distal margin while preserving bowel control and avoiding permanent colostomy. Despite the advantages of these techniques, functional disorders pose a persistent challenge for rectal cancer survivors, with LARS symptoms ranging from daily episodes of incontinence to fecal obstruction and constipation. Presently, there is no targeted therapy specifically for LARS, and management relies on empirical and symptom-based approaches utilizing available treatments for relevant symptoms. Hence, future research endeavors aim to enhance outcomes for cancer patients by delving into the molecular mechanisms underlying LARS and identifying potential therapeutic targets, such as specific genes or pathways, to facilitate more effective interventions. Overall, LARS presents a substantial clinical hurdle, and ongoing efforts strive to enhance patient outcomes and quality of life following rectal cancer surgery. (17),(18),(19)

#### SCRT1

Recent investigations into SCRT1, focusing on the molecular mechanisms influencing radiosensitivity in rectal cancer, unveiled significant findings. Through mRNA data analysis of rectal cancer patients, researchers identified 553 differentially expressed genes (DEGs) out of 600, showcasing high regulation. Among these DEGs, notable downregulated genes were identified, differing significantly between radiation treatments, indicating distinct sensitivities to radiotherapy in rectal cancer. Within these DEGs, five principal genes - TOP2A, MATR3, APOL6, JOSD1, and HOXC6 - emerged as prognostic indicators for rectal cancer through random forest survival analysis. These genes are implicated in various processes, including immune cell invasion, immune-related genes, chemosensitivity, signaling pathways, and transcriptional regulatory networks, underscoring their potential as biomarkers for radiotherapy in rectal cancer. Additionally, while not directly associated with SCRT1, recent studies underscore the significance of lncRNA LINC00974 as a potential therapeutic target. LINC00974 exhibits promise as a clinical predictor and diagnostic tool for tumor growth and metastasis in rectal cancer. In sum, comprehending the roles of genes like TOP2A, MATR3, APOL6, JOSD1, and HOXC6 offers valuable insights into rectal cancer pathogenesis and potential avenues for enhancing radiotherapy outcomes.(20),(21),(22)

#### CEACAM5

Research by Yves Baudat and his team focused on the cancer antigen cell adhesion molecule (CEACAM5), also known as CD66e, a glycosylphosphatidylinositol-anchored glycoprotein prominently expressed on the surface of various epithelial tumors. CEACAM5 is particularly common in almost all colon cancers, with approximately 90% of colon adenocarcinomas showing strong expression, in contrast to limited expression in normal tissue. Studies show that CEACAM5 plays an important role in the pathogenesis of cancer and its escape from the immune system. Key insights into its potential as a target for rectal neoplasms are outlined in research by Chaogu Zheng et al. Anti-CEACAM5 monoclonal antibody (mAb CC4) shows a strong and specific binding affinity to tumor tissues, especially colorectal adenocarcinoma. In xenograft mouse models, mAb CC4 efficiently accumulated at tumor sites and significantly suppressed colorectal tumor growth while enhancing natural killer (NK) cell-mediated tumor immunity. Furthermore, the interaction of mAb CC4 with NK cells enhances their toxicity against MHC-I-deficient colorectal cancer cells and prevents the intercellular interaction between epithelial CEACAM5 and the NK inhibitory receptor CEACAM1, a promising target for preventing tumor immune evasion. Is. These findings demonstrate the potential of mAb CC4 as a novel carrier for tumor targeting and cancer therapy. Overall, CEACAM5 appears as a promising target for the development of innovative cancer therapies, particularly in the field of colorectal neoplasms, with its protective role against NK cell killing underscoring its importance in cancer progression and metastasis.(23),(24)

#### ZNF135

ZNF135, also known as pHZ-17, ZNF61 or ZNF78L1, is a protein involved in colon cancer (CRC), according to studies by Yuan-Hong Xie and colleagues. Its role and potential as a target for rectal neoplasms Colorectal cancer (CRC) is an important global health concern and ranks as the second deadliest cancer and the third most common malignancy worldwide. In 2018, 1.8 million new cases of CRC were reported, resulting in 881,000 deaths, which is approximately 10% of all new cancer cases and deaths worldwide. Despite advances in primary and adjuvant therapies, the prognosis of metastatic CRC remains challenging with a 5-year survival rate of only 12% and targeted therapy for CRC. which has emerged as a promising approach to improve the overall survival of CRC patients. And notable successes include agents such as cetuximab (an anti-EGFR agent) and vaacizumab, an anti-angiogenic agent. And these drugs block specific pathways involved in growth. And CRC progression is effective in prolonging survival. And ZNF135 and colorectal cancer is a protein whose expression has been studied in various cancers including CRC. While the exact role of ZNF135 in CRC is still being elucidated, it is considered a potential target due to its involvement in critical cellular pathways. However, more research is needed to fully understand its mechanisms and therapeutic implications. In addition to limitations and future trends, the role of ZNF135 may become clearer as global guidelines update recommended targeted drugs based on high-quality clinical trials. And researchers are investigating new agents that target different pathways and locations. They block immune checkpoints and aim to improve outcomes for CRC1 patients. Overall, ZNF135 is a potential avenue for targeted therapy in colorectal cancer, but ongoing research will determine its precise impact and clinical significance.(21),(25),(26)

#### ZNF77

Research done by Jing Sun et al. In ZNF77, also known as pHZ-17, ZNF61, or ZNF78L1, its role in colon cancer (CRC) is investigated. CRC is the third most common malignancy worldwide and the second leading cause of cancer-related deaths, with 1.9 million new cases in 2020. ZNF77 expression has been investigated in various cancers including CRC, but its exact role in CRC is still under investigation. However, its association with critical cellular pathways makes it a potential target for CRC therapy, along with other promising targets. Recent research has identified several protein biomarkers associated with CRC risk, and ZNF77 may affect tumorigenic pathways. However, its specific effect and clinical relevance need further investigation. Overall, ZNF77 has potential as a target for CRC therapy, but ongoing research will determine its precise effect and suitability as a therapeutic target.(25),(27),(28)

#### PORCN

PORCN, also identified as pHZ-17, ZNF61, or ZNF78L1, assumes a significant role in colorectal cancer (CRC). The Wnt/β-catenin signaling pathway, crucial for various physiological functions like cell proliferation, differentiation, and tissue balance, is closely intertwined with CRC. Dysregulation of this pathway significantly contributes to the initiation and advancement of solid tumors, including CRC. PORCN serves as a bifunctional regulator, participating in mitosis regulation and programmed cell death inhibition. It notably influences the Wnt/β-catenin pathway by facilitating the production of Wnt ligands, such as Wnt3a, which activates the pathway. Utilization of compounds like LGK974 effectively impedes Wnt3a production by inhibiting PORCN. Preclinical investigations have demonstrated LGK974’s efficacy in reducing tumor growth induced by abnormal Wnt1 expression in four mouse models. Advancing into clinical trials, LGK974 is currently under evaluation, particularly in phase I trials assessing its combination with PDR001, an immune checkpoint inhibitor, for treating BRAF-mutant CRC patients. However, further research is imperative to validate the safety and efficacy of PORCN-targeted agents in CRC treatment. Overall, targeting PORCN within the Wnt/β-catenin pathway holds promise as a prospective therapeutic approach for rectal neoplasms.(29),(30),(31),(32)

#### TRG-GCC3-1

TRG-GCC3-1, also known as pHZ-17 ZNF61 or ZNF78L1, significantly affects tumor regression scores (TRG) in patients undergoing neoadjuvant therapy for rectal cancer, according to research by Jia-yi Li and colleagues. Lays. Understanding the relationship between TRG, pathologic complete response (pCR), and long-term survival is important for evaluating treatment outcomes. A study examining survival prospects and TRG, using reconstructed individual patient data (IPD), examined the relationship between TRG, pCR, and 10-year overall survival (OS) and 5-year disease-free survival (DFS). Contract. Patients who achieved a pCR showed a significant survival rate compared to non-pCR (npCR) patients. Improved tumor regression after neoadjuvant treatment is associated with improved OS and DFS. Notably, patients achieving pCR showed better OS and DFS outcomes compared to patients with npCR. Clinical evidence suggests that increased tumor regression is associated with a better survival outlook, emphasizing the importance of understanding TRG and its implications for patient prognosis in guiding treatment decisions for rectal cancer. Overall, TRG-GCC3-1 appears as a promising pathway for targeted therapy in rectal neoplasms.(9),(12),(25),(33),(34)

#### KRAS

Research conducted by Chongkai Wang and Marwan Fakih delves into KRAS, a prevalent genomic alteration observed in various human cancers, notably playing a crucial role in colorectal cancer (CRC). KRAS Mutations in CRC are detected in roughly 45% of colon cancers, linking these mutations to resistance against EGFR-targeted therapies. RAS-mutated colorectal cancers generally exhibit poorer overall prognoses in contrast to RAS wild-type tumors. Recent strides in KRAS targeting present promising advancements. Specific covalent inhibitors have successfully achieved the knockdown of KRASG12C, a mutation found in 14% of non-small cell lung cancer (NSCLC) cases and 3% of colorectal cancer cases. Noteworthy KRAS-specific inhibitors like sotorasib and MRTX849 have demonstrated encouraging antitumor activity in clinical models and early-phase clinical trials. These inhibitors hold the potential to revolutionize the clinical management of KRAS-mutated solid tumors. Nevertheless, challenges persist in identifying biomarkers predictive of response to KRAS inhibition, and strategies to overcome resistance are actively being explored to optimize clinical benefits. Overall, targeting KRAS presents a promising avenue for enhancing outcomes in rectal neoplasms.(35),(36),(37)

#### CCL20

In studies conducted by Dan Wang and colleagues, CCL20, also referred to as chemokine ligand (C-C motif) 20, emerged as a significant factor influencing chemoresistance in colorectal cancer (CRC) patients undergoing Fulfox chemotherapy, a common treatment regimen. It was observed that resistant patients exhibited higher levels of CCL20 secretion by tumor cells compared to sensitive patients, with elevated CCL20 levels closely linked to chemoresistance and poorer survival outcomes. Particularly, among the drugs used in Fulfox chemotherapy, fluorouracil was noted to increase CCL20 expression in colon cancer cells. The secretion of CCL20 from tumor cells played a crucial role in activating regulatory T cells (Tregs), thereby augmenting chemoresistance to 5-FU. Additionally, the FOXO1/CEBPB/NF-κB signaling pathway was found to be activated in CRC cells following 5-FU treatment, facilitating the upregulation of CCL20. Blocking CCL20 effectively suppressed tumor progression and restored sensitivity to 5-FU in colorectal cancer. The therapeutic potential of targeting the FOXO1/CEBPB/NF-κB/CCL20 axis appears promising for CRC treatment. Inhibiting CCL20 could disrupt Treg recruitment, potentially enhancing chemotherapy efficacy. Targeting this axis holds promise for improving the survival and prognosis of colorectal cancer patients. Overall, comprehending CCL20’s role in promoting chemoresistance and its interaction with the tumor microenvironment offers valuable insights for the development of targeted therapies in rectal neoplasms.(38),(39),(40)

#### SAMM50

Research conducted by Jing Sun and colleagues has delved into SAMM50, a protein showing promise as a potential target in colon cancer. Their study employed proteome-wide analysis, aiming to identify candidate protein markers and therapeutic targets for colorectal cancer (CRC) using proteome-wide Mendelian randomization (MR). Among the proteins scrutinized, SAMM50 emerged as particularly noteworthy. This investigation utilized protein quantitative trait loci (pQTLs) obtained from genome-wide association studies (GWASs) on the plasma proteome. Although SAMM50’s precise role in CRC remains ambiguous, its association with the disease suggests its potential relevance as a therapeutic target. Future research endeavors are necessary to ascertain SAMM50’s viability as a therapeutic intervention target in CRC treatment. Given the significant role of proteomics in identifying drug targets, SAMM50 warrants further exploration for its potential impact on rectal neoplasms. Overall, comprehending SAMM50’s involvement in CRC pathogenesis may offer new insights for devising therapeutic strategies in managing rectal cancer.(41),(42),(26)

#### RAB1A

Research by Menglin Zhu et al. has spotlighted RAB1A, a protein of significant interest, as a prospective target in colorectal cancer (CRC). RAB1A exhibits a close relationship with GLI1 expression, a known driver of CRC metastasis via FoxM1 overexpression. Immunohistochemical analyses unveiled notable upregulation of both RAB1A and FoxM1 in CRC tissues compared to normal tissues. Notably, RAB1A showed a positive correlation with FoxM1 expression, particularly in advanced TNM stages. Elevated expression levels of RAB1A and FoxM1 correlated with unfavorable prognosis in CRC patients. As a member of the RAB family, RAB1A regulates intracellular membrane dynamics, facilitating vesicular trafficking, and influencing signal transduction and cellular autophagy. Prior research has linked RAB1A with oncogenic behavior, including proliferation, migration, and drug resistance across various cancers, suggesting therapeutic implications. Although investigations specific to RAB1A in rectal neoplasms are limited, unraveling its functional role may unveil novel therapeutic avenues. Targeting RAB1A holds promise for impeding CRC progression and enhancing treatment outcomes. However, further exploration is imperative to ascertain its potential as a clinical biomarker and therapeutic target. Overall, RAB1A emerges as a promising candidate for targeted intervention in colorectal cancer, with its interaction with FoxM1 warranting deeper exploration.(43),(44),(45)

#### BRAF

Research conducted by Shuyi Cen and colleagues has focused on BRAF, a pivotal gene within the RAS-RAF-MEK-ERK-MAP kinase pathway, as a potential therapeutic target in colorectal cancer (CRC), encompassing BRAF-mutated rectal neoplasms and their immune microenvironment. Their investigations have revealed distinct features within the tumor microenvironment of patients with BRAF-mutated colon cancer, characterized by heightened stromal cells and augmented immune cell infiltration. Notably, BRAF mutant tumors exhibit elevated levels of stromal cells, fostering increased immune cell penetration. These tumors demonstrate heightened immune cell infiltration but diminished tumor clearance in cases of BRAF mutation. Moreover, several immunotherapy targets, including PD-1, PD-L1, CTLA-4, LAG-3, and TIM-3, are markedly expressed in patients with BRAF mutations. BRAF mutation correlates with specific immune cell subsets characterized by elevated proportions of M1 neutrophils and macrophages and reduced proportions of plasma cells, resting dendritic cells, and naïve CD4 T cells, which exhibit responsiveness to checkpoint inhibitors such as those targeting PD-1/PD-L1. Overall, comprehending the intricate interplay between BRAF mutation and the immune microenvironment holds potential for guiding future treatment strategies for rectal neoplasms.(46),(47),(48),(49)

#### NXN

NXN, also recognized as nucleoredoxin, has been scrutinized concerning rectal neuroendocrine neoplasms (R-NENs) by Mayur Virarkar and colleagues. As a protein implicated in cellular redox regulation, NXN’s precise function in R-NENs remains incompletely understood. Nevertheless, unraveling its role holds potential for shedding light on disease progression and devising therapeutic strategies. Imaging techniques, including endoscopic ultrasound (EUS), computed tomography (CT), and magnetic resonance imaging (MRI), are pivotal in R-NEN diagnosis and management, facilitating the visualization of rectal tumors and enabling early detection for appropriate treatment planning. Therapeutic modalities for R-NEN management hinge on several factors such as tumor stage, size, histologic grade, and lymphatic vessel invasion. Low-risk localized R-NENs may be addressed via endoscopic removal, while high-risk or locally advanced neoplasms may necessitate radical surgery, lymphadenectomy, and/or chemotherapy. Although R-NENs are rare, their incidence has risen due to enhanced screening techniques, prompting ongoing research endeavors aimed at identifying novel targets, including proteins like NXN, for personalized therapeutic approaches. Overall, while specific investigations regarding NXN in R-NEN are limited, continuous research into its function may contribute to enhanced management strategies for rectal neuroendocrine neoplasms. (15), (28),(50),(51)

#### TYMS

Thymidylate synthase (TYMS) has emerged as a potential target in colorectal cancer (CRC) research, as highlighted by Yitong Li et al. In the tumor microenvironment (TME), which is crucial for CRC progression and drug resistance, tumor-associated macrophages (TAMs) play an essential role. TAMs regulate key oncogenic processes in CRC, including tumor initiation, proliferation, metastasis, and drug resistance. M1 macrophages exert tumor suppressive functions, while M2 macrophages promote CRC growth. TAM polarization is influenced by various pathways such as NFKB1, STAT3, WNT5A, and PI3K, and M2 polarization is consistent, controllable, and reversible. Researchers are exploring therapeutic ways to target TAMs to increase treatment efficacy. CRC, a common cancer that causes significant global mortality, is currently managed through approaches such as surgery, chemotherapy, radiotherapy, and targeted therapy. However, drug resistance with resistance mechanisms that include drug degradation, apoptosis inhibition, and DNA damage interference is an important obstacle that affects treatment efficacy. Effective strategies to deal with drug resistance are crucial to improve prognosis. Different cell types and cytokines in the TME shape tumor behavior, leading to new therapeutic interventions for advanced CRC. Efforts to reverse drug resistance by targeting the TME are underway. Overall, TYMS, located in the complex TME, is promising as a potential target for rectal neoplasms. Further research and therapeutic strategies focused on TAMs and TME may provide better outcomes for CRC patients.(33),(52),(53)

#### MSH2

Research conducted by Yi Liu and colleagues in the realm of rectal neoplasms has shed light on the potential therapeutic significance of the MSH2 gene. Radiosensitivity stands as a critical determinant of the efficacy of radiation therapy and patient prognosis in rectal cancer. However, the complete molecular underpinnings governing radiation sensitivity in rectal cancer remain elusive. Recent investigations have delved into radiosensitivity-related genes in rectal cancer through multi-omics methodologies. Among these genes, MSH2 emerges as a notable candidate biomarker for radiotherapy. MSH2 functions as part of the DNA mismatch repair (MMR) system, crucial for preserving genomic stability by rectifying DNA mismatches and averting mutations. Notably, in microsatellite stable colorectal tumors (MSS), alterations in MSH2 correlate with heightened tumor mutational burden (TMB), potentially influencing drug response and overall prognosis in rectal cancer. Additionally, certain miRNAs implicated in drug resistance in colorectal cancer (CRC) directly target MSH2, modulating its expression and function. Understanding these intricate miRNA-MSH2 interactions may pave the way for novel targeted therapies aimed at overcoming drug resistance in CRC. Overall, MSH2 emerges as a promising target for rectal neoplasms, holding potential for therapeutic intervention.(54), (55),(56),(57)

#### MLH1

In research led by Junyong Weng and collaborators within the realm of rectal neoplasms, the MLH1 gene has emerged as a promising avenue for immunotherapy, demonstrating notable therapeutic benefits in the treatment of solid tumors, particularly melanoma and non-small cell lung cancer (NSCLC). Immunotherapy, specifically immune checkpoint blockade (ICB), has yielded enduring advantages for select patient cohorts. Regrettably, the efficacy of immunotherapy remains largely confined to a subset of colorectal cancer (CRC) patients exhibiting microsatellite instability (MSI-H). MSI-H tumors exhibit deficiencies in DNA mismatch repair (MMR) genes, including MLH1. Mutations in MLH1 impact tumor mutational burden (TMB) and drug responsiveness in CRC. Ongoing research endeavors aim to identify safety checkpoints to sustain therapeutic effects in patients, particularly those with metastatic CRC (mCRC). MLH1 plays a pivotal role in the MMR system, essential for maintaining genomic integrity. Consequently, its hypermethylation status and expression levels hold promise as predictive biomarkers for the efficacy of immune checkpoint blockade. Clinical trials are actively exploring the impact of immune checkpoints with the objective of enhancing outcomes in CRC. Overall, MLH1 emerges as a compelling target for rectal neoplasms with potential therapeutic implications.(58),(59),(60),(61)

#### DPYD

Research conducted by Ying Yang and colleagues focusing on the DPYD gene has underscored its potential significance in treatment approaches. DPYD encodes the dihydropyrimidine dehydrogenase enzyme, which holds a pivotal role in the metabolism of fluoropyrimidine-based chemotherapy drugs commonly utilized in colorectal cancer (CRC) treatment. In recent times, several investigations have identified microRNAs (miRNAs) implicated in regulating drug resistance in CRC. Certain miRNAs have surfaced as promising targets for mitigating drug resistance in CRC. Preemptive genotyping for DPYD has been proposed as a cost-effective strategy prior to fluoropyrimidine-based chemotherapy. Overall, comprehending the involvement of DPYD and its interplay with miRNAs promises valuable insights for advancing targeted therapeutic modalities in rectal neoplasms management.(56),(62),(63)

#### OR10H3

Research conducted by Al B. Benson and colleagues has brought the OR10H3 gene to the forefront as a prospective target. Although its specific role remains to be fully elucidated, it garners attention in the NCCN guidelines for rectal cancer, particularly in the context of managing malignant polyps and resectable nonmetastatic rectal cancer. Notably, complete neoadjuvant therapy (TNT) assumes a significant role across disease stages. A novel approach termed “watch and wait” presents a non-surgical management option for patients exhibiting a complete response to neoadjuvant therapy. The comprehensive NCCN guidelines encompass various facets such as risk assessment, pathology, staging, management of metastatic disease, post-treatment monitoring, handling of recurrent diseases, and survival outcomes in colorectal cancer (CRC). Despite CRC posing a substantial health challenge, advancements in screening, early detection, and treatment have contributed to declines in its incidence and mortality rates. However, racial disparities persist, with elevated rates in non-Hispanic blacks and lower rates in Asian/Pacific Islander Americans. Furthermore, there is a concerning trend of increasing incidence among individuals younger than 65 years, underscoring the importance of early detection efforts. TNT emerges as a promising approach in rectal cancer management, involving comprehensive neoadjuvant treatment preceding surgery, with ongoing comparative studies against conventional chemotherapy (CRT) to assess its efficacy. While further investigation is warranted regarding the specific role of OR10H3 in rectal neoplasms, these guidelines and ongoing research endeavors provide valuable insights into the evolving landscape of rectal cancer management.(63),(64),(65)

#### OR6J1

Research led by Al B. Benson and collaborators has highlighted the potential significance of the OR6J1 gene as a therapeutic target. This observation is reflected in the NCCN Clinical Practice Guidelines for Rectal Cancer (2022), which extensively addresses the management of malignant polyps and resectable nonmetastatic rectal cancer. The guidelines have been updated to include recommendations pertaining to complete neoadjuvant therapy, radiation modalities, and a non-surgical “watch and wait” strategy tailored for patients exhibiting a complete response to neoadjuvant therapy. Furthermore, the American Association of Colon and Rectal Surgeons (2022) Clinical Practice Guidelines offer comprehensive coverage on the diagnosis, staging, treatment, and post-treatment monitoring of colon cancer patients. While not specifically targeting OR6J1, these guidelines offer valuable insights into rectal cancer management. Notably, a recent study presented at ASCO 2022 reported a notable 100% complete response rate in patients diagnosed with mismatch repair-deficient (MMRd) rectal cancer, particularly in locally advanced cases treated with immunoablative neoadjuvant immunotherapy. Although this study does not directly delve into OR6J1, it underscores promising strides in rectal cancer treatment.(66),(67) Recent investigations led by Hannah Shailes and her team have underscored the Adenomatous Polyposis Coli (APC) tumor suppressor gene as a viable target in the management of rectal neoplasms. Mutations within the APC gene are prevalent, occurring in approximately 80% of sporadic colorectal cancer (CRC) cases, and are also implicated in familial adenomatous polyposis (FAP), an inherited form of CRC. APC functions as a crucial antagonist within the Wnt signaling pathway by downregulating β-catenin, a key player in cell proliferation regulation. Notably, researchers have unveiled a novel synthetic lethal relationship between APC mutations and a class of drugs known as HMG-CoA Reductase inhibitors, commonly referred to as statins. Statins induce a marked reduction in cell viability specifically in APC mutant cell lines compared to their APC wild-type counterparts. Mechanistically, this interaction results in diminished Wnt signaling and reduced expression of the anti-apoptotic protein Survivin, particularly in APC mutant cells, via Rac1-mediated control of β-catenin. The discovery of this synthetic lethal association presents promising avenues for targeted cancer therapy, with statins—already FDA-approved for other indications—potentially serving as a treatment option for APC-mutated CRC. Overall, comprehending the intricate interplay involving APC and its downstream effects on Wnt signaling holds significant promise for advancing targeted therapies against rectal neoplasms. Further exploration through research and clinical trials is imperative to validate these findings and translate them into efficacious treatments.(68),(69),(70)

#### TP53

Lijun Du and colleagues have investigated the pivotal role of the TP53 gene and its potential as a therapeutic target. TP53 is notably implicated in inflammatory bowel disease-associated colorectal cancer (IBD-CRC), with research suggesting that TP53 mutations could serve as valuable biomarkers for screening cancer and dysplasia in individuals with inflammatory bowel disease (IBD). These mutations present a targetable avenue for intervention. Michael P. East and colleagues have identified PIP5K1A, a gene integral to cell signaling, as another potential target for cancers harboring KRAS or TP53 mutations. Loss of PIP5K1A leads to the downregulation of downstream markers associated with oncogenic KRAS signaling and cell proliferation, particularly in the context of TP53 mutation. Francesco Sclafani et al. While specific randomized trials evaluating anti-EGFR-based strategies in resectable rectal cancer are lacking, analyzing TP53 mutations in randomized controlled trials for adjuvant colon cancer and metastatic colorectal cancer could yield valuable insights.

Overall, comprehending the significance of TP53 mutations and their implications for rectal neoplasms may inform targeted therapeutic strategies and bolster treatment approaches for patients with colorectal cancer.(71), (72)

#### NRAS

Kang Chen and colleagues have explored the NRAS gene, a crucial player in cell signaling pathways, implicated in various cancers, including colorectal cancer (CRC). However, as of 2022, there are no approved therapies directly targeting NRAS mutations in CRC. These mutations are observed in about 19% of cancers and exert significant influence on tumorigenesis and tumor progression. The predominant NRAS mutations, namely G12, G13, and Q61, constitute approximately 96% to 98% of all NRAS isoform mutations. Patients diagnosed with NRAS-mutated colorectal cancer typically undergo conventional chemotherapy regimens like FOLFOX, FOLFIRI, or CAPOX, sometimes without the addition of bevacizumab. Although NRAS mutations are commonly associated with CRC, the pursuit of targeted therapies tailored to address NRAS mutations remains an ongoing area of research and development.(73),(74)

#### TYMP

In a study conducted in 2022 by Ankush Paladhi and his team, the focus was on investigating the potential of thymidine phosphorylase (TYMP) as a therapeutic target for rectal neoplasms. The research delved into the connection between T-cell exhaustion and TYMP, which plays a crucial role in conferring resistance to immunotherapy in microsatellite-resistant colorectal cancer (CRC). Their findings revealed that TYMP plays a significant role in triggering systemic T-cell exhaustion and undermining treatment effectiveness alongside dendritic cells (DC) in the CRC model. To target TYMP, the researchers utilized tipiracil hydrochloride (TPI). Treatment with TPI induced immunological cell death (ICD), and the combined approach of TPI and imiquimod-activated DCs led to the induction of ICD in vivo, transforming CRC tumors into immunologically “warm” tumors. High-dimensional cytometric analysis indicated a dependency on treatment outcome concerning T cells and IFN-γ. Moreover, chemoimmunotherapy resulted in the conversion of intratumoral Treg cells into effector Th1 cells, elimination of tumor-associated macrophages, and enhanced T-lymphocyte infiltration and activation, thereby increasing cytotoxicity. This effect was linked to the reduction of PD-L1 expression in tumors and the prevention of T-cell exhaustion. The synergistic interactions between dendritic cells and TPI-induced immunogenic cell death amplified the immune response and tumoricidal activities against microsatellite-resistant colorectal cancer. Targeting TYMP holds promise in augmenting the effects and outcomes of DC immunotherapy in CRC.(75),(76),(77),(78)

#### PIK3CA

The research conducted by Ioannis A. Voutsadakis on the PIK3CA gene as a potential treatment target focused on PIK3CA mutations in colorectal cancer (CRC), which code for the catalytic subunit of the PI3K kinase. These mutations are prevalent in both colorectal cancer cell lines and patient samples, suggesting their significance in CRC tumorigenesis through the activation of the PI3K/AKT/mTOR signaling pathway. To assess this, the researchers examined a range of colon cancer cell lines, comparing those with PIK3CA mutations to those without. They found that cell lines with PIK3CA mutations exhibited varying degrees of sensitivity to certain PI3K inhibitors, although not all. Notably, microsatellite unstable cell lines (MSI) and microsatellite stable cell lines (MSS) demonstrated heightened sensitivity to PI3K inhibitors. Additionally, they observed concurrent mutations in the PI3K/AKT and KRAS/BRAF/MEK/ERK pathways in the sensitive cell lines. The differing susceptibilities among PIK3CA mutant cell lines suggest the involvement of additional molecular abnormalities in determining sensitivity. Consequently, targeting the PI3K pathway, especially in the context of PIK3CA mutations, shows promise for managing rectal neoplasms and enhancing treatment outcomes.(79),(80),(81)

#### CDX2

Research conducted by Kristine Aasebø and colleagues on the CDX2 gene, a member of the tail-type homeobox family, has shed light on its role as an intestinal-specific transcription factor crucial for intestinal growth and differentiation. CDX2 expression holds significant prognostic and predictive value in metastatic colorectal cancer (mCRC). Reduced CDX2 expression correlates with poorer differentiation, microsatellite instability, and specific mutations such as BRAF and KRAS. Patients lacking CDX2 expression tend to receive less chemotherapy and exhibit inferior outcomes. Moreover, rapid progression to first-line combination chemotherapy is more prevalent among CDX2-deficient patients. Median overall survival (OS) is notably shorter in cases where CDX2 expression is lost compared to those with intact CDX2 expression. Recent research suggests that CDX2 restoration therapy may serve as a potential therapeutic approach, with low CDX2 status significantly influencing treatment effectiveness. A 50-gene signature indicative of treatment response indicates that CDX2 reinstatement therapy could potentially halve the risk of death/recurrence. Although CDX2 mutations are rare in colorectal cancer (CRC), the loss of CDX2 expression is frequently observed, particularly in unstable right-sided and microsatellite tumors, where it is associated with aggressive tumor behavior. Overall, CDX2 presents itself as a promising target for rectal neoplasms, offering potential both as a prognostic marker and as a therapeutic intervention. (81),(82),(83)

#### KRT20

Research conducted by Ji Ae Lee and colleagues on the KRT20 gene reveals an intriguing aspect of its function. This gene serves as one of the distinctive markers of the intestinal epithelium commonly utilized in immunohistochemical analyses. Such utilization aids in the identification of cells possessing characteristic features of the intestinal lining. Notably, certain colorectal carcinomas (CRC) exhibit diminished expression of KRT20, alongside other markers like CDX2 (homeobox tail type 2) and SATB2 (specific AT-rich sequence-binding protein 2). These reductions are particularly noticeable in microsatellite-high instability (MSI-H) systems. The diminished presence of these markers suggests that KRT20 could serve as a viable target for therapeutic intervention in rectal neoplasms. Further comprehension of its role and regulatory mechanisms holds the potential to pave the way for novel therapeutic approaches.(59),(63),(84)

#### CCNO

Research conducted by Al B. Benson and colleagues sheds light on the CCNO (Cyclin O) gene, a regulator of cyclin-dependent kinase involved in cell cycle advancement. This gene plays a pivotal role in facilitating the transition from G1 to S phase, thereby fostering cell proliferation. Recent investigations have underscored CCNO as a prospective target in cancer therapy, given its propensity for overexpression across various cancer types, including colon cancer. In the context of rectal neoplasms, CCNO’s heightened expression may fuel unbridled cell division and tumor expansion. Consequently, targeting CCNO holds promise in thwarting cell cycle progression and curbing tumor proliferation. Nonetheless, it is imperative to acknowledge the necessity for further studies to comprehensively elucidate the precise mechanisms and therapeutic ramifications of CCNO in rectal cancer. As clinical trials and additional research endeavors unfold, they are poised to furnish invaluable insights into CCNO’s potential as a therapeutic target. It is imperative to bear in mind that scientific advancements are ongoing, and researchers persist in unearthing novel modalities for rectal neoplasm treatment..(15),(64)

### 1.2 Rectal Neoplasm treatment

Targeted Therapy:

#### Bevacizumab

Bevacizumab represents a crucial component in the therapeutic arsenal against metastatic colorectal cancer (mCRC). As a targeted therapy falling under the class of angiogenesis inhibitors, bevacizumab exerts its effects by selectively binding to vascular endothelial growth factor (VEGF), impeding its interaction with cell surface receptors. This action primarily curtails the microvascular proliferation of tumor blood vessels, thereby constraining the blood supply to tumor tissues and impeding their proliferation. Typically, bevacizumab is administered alongside other chemotherapy agents. Neoadjuvant therapy is administered preoperatively to shrink tumor size and enhance surgical outcomes, while adjuvant treatment follows surgery to forestall recurrence. In advanced cases, palliative therapy aims to alleviate symptoms and extend life expectancy. A systematic review and meta-analysis of randomized controlled trials (RCTs) revealed that bevacizumab enhances antitumor activity and disease management, particularly as a monotherapy. When combined with other agents like cetuximab and panitumumab, it augments disease control benefits, albeit without significant enhancements in overall survival or progression-free survival. Being a biotechnology drug, bevacizumab’s capacity to inhibit angiogenesis underscores its value in colorectal cancer treatment. Nevertheless, personalized treatment strategies should be tailored to individual patient characteristics and informed by clinical evidence.(85), (86)

#### Triiodothyronine

Triiodothyronine (T3), a thyroid hormone, assumes a crucial role in regulating diverse physiological functions. Synthesized by the thyroid gland, it is indispensable for normal growth, neural differentiation, and metabolic control in mammals. T3 binds to thyroid hormone receptors (TR), nuclear receptors facilitating gene expression modulation upon entering target cells through membrane iodothyronine transporters. Its interaction with specific TR isoforms and coactivators/chronorepressors elicits transcriptional changes impacting cellular functions. Although not conventionally employed as a direct therapy for rectal neoplasms, the metabolic and cellular effects of T3 are noteworthy. It influences energy expenditure, lipid metabolism, and glucose utilization, offering potential implications for managing obesity and hyperlipidemia. Preliminary studies suggest a role in enhancing cognitive function, albeit requiring further investigation. Ongoing research explores its implications in heart health and potential synergy with other cancer treatment modalities, such as chemotherapy and immunotherapy, to bolster overall efficacy. While T3 itself isn’t a standalone therapy for rectal neoplasms, comprehending its mechanisms and potential applications may inform future therapeutic approaches.(87), (88)

#### Thyroxine

Thyroxine (T4), also referred to as levothyroxine, is an artificially produced variant of the thyroid hormone generated by the thyroid gland. Serving as a primary endogenous hormone, T4 holds a crucial role in regulating metabolism, growth, and developmental processes. In instances like hypothyroidism, where the thyroid gland inadequately produces thyroid hormone, T4 serves as a replacement. Through enzymatic deiodination by either type I or type II 5’-deiodinase, T3, its active metabolite, is derived from T4. The physiological effects of T3 encompass a wide range, mirroring those of thyroid hormones, including metabolic regulation and modulation of heart function and gene expression. Levothyroxine is frequently administered for hypothyroidism treatment as part of an alternative therapeutic approach. By reinstating thyroid hormone levels, it alleviates symptoms like fatigue, weight gain, and intolerance to cold. Tailoring the dosage to individual patient needs and thyroid-stimulating hormone (TSH) serum levels is imperative. While T4 itself isn’t directly utilized as targeted therapy or immunotherapy for rectal neoplasms, ensuring optimal thyroid function throughout cancer treatment holds paramount importance. T4 levels can impact overall health, immune response, and treatment efficacy.(89),(90)

#### Deoxycytidine

Beverages containing dietary supplements are designed to offer extra nutrients, vitamins, and minerals. They are commonly utilized to bolster general health, particularly in individuals battling cancer, who often require additional nutrition due to the illness and its therapies. While dietary supplement drinks are typically not regarded as the primary treatment for rectal neoplasms, they play a crucial supportive role in overall cancer management. These supplements can complement standard treatments like surgery, chemotherapy, or radiation therapy. Cancer patients frequently encounter weight loss, muscle depletion, and malnutrition, and nutritional supplements aim to prevent or alleviate these effects. Adequate nutrition is vital for supporting the immune system, particularly for cancer patients. Nutritional supplements can help boost energy levels, stimulate appetite, and improve overall well-being by providing essential nutrients, including vitamins (e.g., vitamin D), minerals (e.g., zinc), and amino acids (e.g., glutamine), which are crucial for various cellular processes and immune function. Biotech nutraceuticals encompass specialized formulations containing proteins, peptides, or other bioactive compounds, such as immune-enhancing formulas or specific amino acid blends. These supplements support cellular metabolism, tissue repair, and immune system function. For instance, glutamine has been investigated for its potential to mitigate inflammation and enhance clinical outcomes in colorectal cancer patients. Recent studies have explored the impact of specific nutritional supplements on inflammation, nutritional status, and clinical results in colorectal cancer patients. Veglutamine, for example, has shown promise in reducing inflammatory markers and shortening hospital stays. However, additional randomized controlled trials (RCTs) are needed to validate these findings and explore other nutritional supplements further. Overall, while dietary supplement beverages are not standalone remedies for rectal neoplasms, they serve as complementary measures to promote overall health and well-being during cancer treatment.(91)

### Chemotherapy

#### Fluorouracil

Fluorouracil (5-FU) is a frequently utilized chemotherapy medication for managing rectal neoplasms (colorectal cancer). It falls into the class of anti-metabolites and functions by primarily inhibiting thymidylate synthase (TS), a crucial enzyme necessary for DNA synthesis. By blocking TS activity, 5-FU disrupts the production of thymidine, which hampers DNA replication and induces cell death. Typically, 5-FU is administered alongside other chemotherapy drugs like oxaliplatin and irinotecan to enhance treatment efficacy by targeting various aspects of cancer cell proliferation and survival. Occasionally, targeted therapies like cetuximab (an anti-EGFR antibody) are combined with 5-FU to hinder cancer cell signaling pathways. The metabolite 5-FUTP, which is extensively integrated into RNA, interferes with regular RNA processing and function, resulting in mis-splicing that can provoke RNA toxicity at multiple levels, impacting cancer cell viability. Efforts to surmount 5-FU resistance persist, with researchers exploring novel strategies employing advanced technologies like CRISPR-Cas, TALENS, and patient-derived xenograft models to refine clinical relevance and precision. Overall, fluorouracil remains pivotal in colorectal cancer treatment, and ongoing research endeavors aim to optimize its utilization and address resistance issues.(92),(93),(73)

#### Capecitabine

Capecitabine is a chemotherapy medication frequently employed in rectal neoplasm treatment. It falls within the class of anti-metabolites, acting by disrupting the production of DNA, RNA, and proteins in the body. This interference halts or slows the growth of cancer cells and other rapidly dividing cells, ultimately leading to their demise. Clinically, capecitabine is often utilized alongside other chemotherapy agents like oxaliplatin and irinotecan to enhance treatment efficacy by targeting distinct facets of cancer cell growth and survival. Additionally, targeted therapies such as cetuximab (an anti-EGFR antibody) may be combined with capecitabine to impede cancer cell signaling pathways. Capecitabine is employed for specific applications, including colon cancer treatment alone or in conjunction with other chemotherapy following surgery to prevent cancer recurrence. It is also administered around the time of surgery, along with radiation and other chemotherapy, in adults with rectal cancer that has spread to nearby tissues. Furthermore, capecitabine is prescribed alone or with other chemotherapy agents in cases where colorectal cancer is inoperable or has metastasized. It finds utility in pancreatic adenocarcinoma as part of post-surgical therapy to prevent cancer recurrence. Moreover, capecitabine is used in esophageal cancer alongside other chemotherapy drugs when surgery is not feasible or when the cancer has spread. In HER2-positive gastric cancer, it is employed in cases where the cancer has spread and has not been treated with a chemotherapy regimen involving capecitabine. Ongoing research endeavors strive to optimize capecitabine usage and address resistance issues, with technologies such as CRISPR-Cas, TALENS, and patient-derived xenograft models aiding in treatment refinement. Overall, capecitabine remains a crucial element in colorectal cancer treatment, and ongoing research endeavors aim to maximize its therapeutic effectiveness.(94),(95)

#### Oxaliplatin

Oxaliplatin is a chemotherapy medication typically employed in treating colorectal cancer, including rectal neoplasms. It belongs to the class of platinum-based chemotherapy drugs and operates by causing DNA damage within cancer cells. Specifically, oxaliplatin forms covalent bonds with macromolecules, disrupting DNA replication and transcription processes, ultimately resulting in cell death. Typically, oxaliplatin is administered alongside other chemotherapy drugs such as fluorouracil (FU) and leucovorin, constituting the FOLFOX regimen, a systemic chemotherapy utilized for colorectal cancer treatment, including rectal neoplasms. Unlike targeted therapy or immunotherapy, oxaliplatin falls within the realm of conventional cytotoxic chemotherapy and is a small molecule chemotherapy agent rather than a biological or biotechnological medicine. A study highlighted in PMC Europe demonstrates that oxaliplatin hinders colorectal cancer progression by targeting the chemokine CXCL11 secreted by cancer-associated fibroblasts, impeding progression through the CXCR3/PI3K/AKT pathway. Overall, oxaliplatin significantly contributes to colon cancer treatment, including rectal neoplasms, given its DNA-disrupting mechanism leading to cell death. However, tailoring treatment approaches based on individual patient factors and consulting with oncologists remain crucial for determining the most suitable course of action.(96),(97)

#### Modified FOLFOX6 Regimen

The modified FOLFOX6 regimen (mFOLFOX6) stands as a chemotherapy protocol widely employed in managing colorectal cancer, including rectal neoplasms. Its mechanism of action combines several drugs: oxaliplatin, a platinum-based chemotherapy agent; fluorouracil (5-FU), a fluoropyrimidine antimetabolite; and calcium folinate (Leucovorin), a folate analog. Oxaliplatin forges covalent bonds with DNA, disrupting replication and transcription processes within cancer cells, ultimately inducing cell death. Fluorouracil, meanwhile, inhibits thymidylate synthase, pivotal for DNA synthesis, further impeding cancer cell growth. Administered cyclically intravenously, mFOLFOX6 does not fall under targeted therapy or immunotherapy but rather conventional cytotoxic chemotherapy. Its adoption is justified by comparable efficacy to the standard FOLFOX4 regimen, sans the need for a fluorouracil bolus on day 3. Although head-to-head randomized controlled trials (RCTs) directly contrasting mFOLFOX4 with mFOLFOX6 in the adjuvant setting are lacking, mFOLFOX6 remains a valuable option for rectal neoplasm treatment. However, selecting the most effective treatment method necessitates consideration of individual patient factors and consultation with oncologists.(98),(99) The Modified CAPOX (mCAPOX) regimen stands as a chemotherapy combination utilized in colorectal cancer treatment, encompassing rectal neoplasms. Its mechanism involves two key drugs: Capecitabine, an oral prodrug converting to 5-fluorouracil (5-FU), and Oxaliplatin, a platinum-based chemotherapy agent. Capecitabine hinders thymidylate synthase, vital for DNA synthesis, disrupting cancer cell growth. Oxaliplatin forges covalent DNA bonds, impeding replication and transcription processes, ultimately prompting cell demise. Administered in a 21-day cycle, oxaliplatin intravenously over two hours on cycle day one, while capecitabine is orally taken twice daily from day one’s evening to day fifteen’s morning. Total daily dose ranges from 1700 to 2000 mg/m2. mCAPOX isn’t targeted therapy or immunotherapy but falls under conventional cytotoxic chemotherapy. Its aim is to optimize effectiveness. Both capecitabine and oxaliplatin are small molecule agents. A JAMA Network Open study explored bevacizumab’s combination with standard oxaliplatin-based regimens, including mCAPOX, scrutinizing diverse prescribing schedules and their impact on colorectal cancer treatment. Overall, mCAPOX presents a valuable rectal neoplasm treatment option, yet individual patient factors and oncologist consultation guide effective treatment selection.(86)

#### Oxaliplatin

The Modified Long-Course Chemotherapy Regimen (mLCRT) represents a holistic strategy for rectal neoplasms, integrating chemotherapy and radiation therapy. mLCRT’s mechanism entails an extended course of chemotherapy pre-surgery, aiming to reduce tumor size, enhancing surgical efficacy. Combining radiotherapy with chemotherapy bolsters local tumor control and curbs distant metastases’ risk. While mLCRT doesn’t specify particular drugs, it typically involves fluoropyrimidine-based regimens (like 5-FU or capecitabine) coupled with radiation. Administered over around 5.5 weeks, mLCRT isn’t targeted therapy or immunotherapy but falls within conventional cytotoxic chemotherapy with radiation. The objective is tumor shrinkage, bolstered local control, and enabling sphincter-preserving surgery. Total neoadjuvant strategies, encompassing mLCRT, amalgamating prior chemotherapy and radiotherapy, seek to optimize outcomes. 5-FU or capecitabine, utilized in chemotherapy, are small molecule drugs, whereas radiation therapy is non-pharmacological. An investigation in BMC Cancer underscores the necessity of a specialized multidisciplinary approach to achieve satisfactory tumor regression and local recurrence control in rectal cancer patients on neoadjuvant chemotherapy. Overall, mLCRT substantially enhances surgical outcomes for rectal neoplasms, yet personalized patient factors and oncologist input are pivotal in method selection.(22), (100), (101)

#### FOLFOX Regimen

The FOLFOX regimen, a chemotherapy protocol mainly for colorectal cancer treatment, comprises three drugs: folinic acid (FOL), oxaliplatin (OX), and fluorouracil (5-FU). FOLFOX4 adjuvant therapy, recommended for colon cancer patients, involves 12 cycles every two weeks. On day 1, intravenous (IV) infusion of oxaliplatin (85 mg/m2) and simultaneous IV infusion of leucovorin (200 mg/m^²^) occur, along with 5-FU (400 mg/m2) IV bolus followed by a 22-hour 5-FU (600 mg/m2) IV infusion. On day 2, leucovorin (200 mg/m^²^) IV infusion precedes a 5-FU (400 mg/m^²^) IV bolus and another 22-hour IV infusion of 5-FU (600 mg/m2). FOLFOX6 offers an alternate dosing schedule. Among the components, fluorouracil (5-FU) and oxaliplatin are small molecule drugs, while folinic acid (Leucovorin) is a biotechnology drug due to its specific cancer treatment utility, although derived from natural folates. Oxaliplatin disrupts DNA replication and transcription via DNA cross-links, prompting cancer cell apoptosis, while 5-FU impedes thymidylate synthase and diminishes DNA synthesis, also affecting RNA function. Leucovorin bolsters 5-FU’s efficacy by stabilizing its binding to thymidylate synthase. Overall, the FOLFOX regimen’s combination of these drugs targets cancer cells via diverse mechanisms, ultimately enhancing treatment outcomes for patients recovering from rectal neoplasms.(102),(103),(104)

#### Semustine

Recent literature highlights semustine, also referred to as methyl-CCNU, as an alkylating chemotherapy agent categorized under small molecule drugs. Its primary application lies in treating various cancers, including rectal neoplasms. Semustine, a member of the nitrosoureas family, exerts its action by alkylating DNA and can be administered orally or intravenously, either alone or in conjunction with other chemotherapy medications.

Although less common in rectal neoplasms, vesmustine is utilized in brain tumors, Hodgkin’s lymphoma, and other malignancies, albeit selectively. The mechanism of action involves DNA alkylation, where semustine forms covalent DNA bonds, inducing cross-linkage between DNA strands. This interference impedes DNA replication and transcription, ultimately impeding cancer cell growth and division. Moreover, vesmustine triggers apoptosis in cancer cells, thus curbing their uncontrolled proliferation. Its anti-tumor effect primarily targets rapidly dividing cells, encompassing cancer cells, influencing both tumor initiation and progression. However, vesmustine may suppress bone marrow, leading to diminished blood cell production, alongside common side effects like nausea, vomiting, and fatigue. Regular monitoring and supportive care during treatment are indispensable. In essence, semustine, functioning as a small molecule alkylating agent, disrupts cancer cell DNA, impeding their growth. Although not a front-line therapy for rectal neoplasms, it might be a viable consideration in specific clinical contexts.(105)

#### FOLFIRI Regimen

FOLFIRI, a chemotherapy protocol employed in colorectal cancer management, consists of three key components: FOL, denoting calcium leucovorin, which is a form of folinic acid, a derivative of vitamin B. Leucovorin enhances the cytotoxic effects of 5-fluorouracil (5-FU). The “F” represents fluorouracil, an antimetabolite that hampers DNA synthesis. “IRI” refers to irinotecan hydrochloride, a topoisomerase I inhibitor disrupting DNA replication. Leucovorin’s mechanism of action involves augmenting 5-FU activity by bolstering its binding to thymidylate synthase (TS), thereby impeding DNA synthesis. Fluorouracil binds with RNA and DNA, leading to cell cycle arrest and subsequent cell demise. Irinotecan suppresses topoisomerase, causing DNA damage and inducing apoptosis in cancer cells. FOLFIRI demonstrates efficacy in treating metastatic colorectal and gastric cancers, but its effectiveness in adjuvant colon and rectal cancer treatment remains unsubstantiated. FOLFIRI comprises small molecule agents, including leucovorin, fluorouracil, and irinotecan. The unique mechanisms of action of these drugs collectively render FOLFIRI a valuable therapeutic option for colon and gastric cancers. Nonetheless, its utility in adjuvant therapy for colon and rectal cancer is constrained.(106)

#### Preoperative Chemotherapy

Recent studies have highlighted the objective of preoperative chemotherapy as reducing tumor burden before surgery, aiming to eradicate microscopic cancer cells that may elude detection and potentially metastasize. The primary aim is curbing cancer recurrence post-surgery. Preoperative chemotherapy encompasses neoadjuvant chemotherapy administered prior to surgery to diminish tumor size and adjuvant chemotherapy given post-surgery to lower recurrence risks. It is also utilized for palliative care, enhancing the quality and prolonging life when curative treatment is unfeasible. Preoperative chemotherapy employs both small molecule drugs and biotechnology agents, with specific drug selection tailored to individual patient factors and treatment plans. These drugs exert diverse mechanisms: cytotoxic effects directly target cancer cells, impeding their growth and division, cell cycle inhibition hampers cell proliferation, DNA damage-inducing drugs prompt cell death, while angiogenesis inhibitors thwart tumor blood vessel formation. Certain biotechnology drugs diminish the immune system’s capability to recognize and combat cancer cells. Notably, a significant study compared neoadjuvant FOLFOX (fluorouracil, leucovorin, and oxaliplatin) with standard chemotherapy for locally advanced rectal cancer, demonstrating non-inferiority in disease-free survival rates with neoadjuvant FOLFOX compared to chemoradiotherapy. The five-year disease-free survival rate was approximately 80.8% in the FOLFOX group and 78.6% in the chemotherapy group. Overall, preoperative chemotherapy, including regimens like FOLFOX, plays a pivotal role in rectal neoplasm management.(107)

#### FOLFOXIRI Regimen

FOLFOXIRI is a blend of chemotherapy medications employed in the management of advanced or metastatic colorectal cancer. Consisting of Fluorouracil (5FU), oxaliplatin, and irinotecan, these agents target rapidly dividing cells, including cancerous ones, as they circulate throughout the bloodstream. While folinic acid is not itself a chemotherapy drug, it is administered alongside fluorouracil to enhance its efficacy. The FOLFOXIRI regimen combines small molecule drugs (fluorouracil, oxaliplatin, and irinotecan) with biological components (folinic acid), delivered via intravenous infusion. It is utilized as standalone therapy or in conjunction with platinum agents and radiation therapy, or with leucovorin, fluorouracil, and irinotecan chemotherapy. FOLFOXIRI is frequently integrated with other treatments such as targeted therapies or immunotherapy, with treatment decisions tailored to individual patient characteristics, tumor attributes, and treatment response.(106)

### Antiviral Medication

#### Irinotecan

Irinotecan, an antineoplastic enzyme inhibitor, is predominantly employed in colorectal cancer therapy, originating from camptocin. Its mechanism involves blocking topoisomerase, pivotal in DNA replication and repair. Irinotecan binds to the I-DNA topoisomerase complex, hindering DNA strand reconnection, thereby causing double-stranded DNA breaks and eventual cell demise. By forming a ternary complex with topoisomerase I and DNA, it obstructs the mobile replication fork during DNA replication, halting replication and inducing fatal double-strand DNA breaks. Essentially, Irinotecan disrupts cancer cell division by impeding topoisomerase, crucial for DNA unwinding and repair. Effective in metastatic colorectal cancer, small cell lung cancer, and metastatic or recurrent cervical cancer, as well as metastatic adenocarcinoma of the pancreas, irinotecan is a small molecule drug. A study revealed that normal irinotecan doses reduce tumor oxygenation and blood perfusion via an anti-angiogenic mechanism, corroborated by immunohistochemistry assessing hypoxia and endothelial cell marker CD312. Overall, irinotecan’s topoisomerase inhibition renders it valuable in rectal neoplasm treatment, particularly when combined with other agents.(86)

#### Tegafur

Tegafur, categorized as an antineoplastic agent among pyrimidine analogs, is utilized alongside other anticancer medications to combat advanced gastric and colorectal cancers. Functioning as a prodrug of fluorouracil (5-FU), tegafur metabolizes into 5-FU within the body. Upon conversion, it obstructs thymidylate synthase (TS) along the pyrimidine pathway involved in DNA synthesis. By impeding the production of 2’-deoxythymidylate (DTMP), tegafur effectively halts DNA synthesis, crucial for cancer cell growth and proliferation. Typically combined with other drugs to enhance 5-FU’s bioavailability or mitigate toxicity, tegafur is prescribed in various combinations for different cancers. For instance, tegafur-uracil, together with fluorouracil, treats colorectal cancer, while tegafur, gimeracil, and oteracil (S-1) address advanced gastric cancer. Additionally, in stomach cancer, tegafur with cisplatin is utilized for advanced cases. As a small molecule, tegafur, when coupled with uracil and calcium folinate, serves as the primary treatment for metastatic colorectal cancer. Overall, tegafur’s disruption of DNA synthesis renders it a valuable weapon against neoplasms, with combination therapies and targeted mechanisms enhancing its efficacy in advanced gastric and colorectal cancers.(108)

#### Mitomycin

Mitomycin, an anticancer antibiotic originally discovered by Japanese microbiologists in the 1950s from Streptomyces caespitosus cultures, functions as an alkylating agent, uniquely inhibiting DNA synthesis by forming cross-links between the DNA’s complementary strands. This alkylating mechanism sets mitomycin apart from most other antibiotics. When activated in vivo, mitomycin acts as both a bifunctional and trifunctional alkylating agent, binding to DNA to disrupt replication and transcription processes. At higher concentrations, it extends its impact to RNA and protein synthesis. Mitomycin’s efficacy has warranted its approval for various cancers, including low-grade upper urinary tract cancer (LG-UTUC), and it is additionally employed as an adjunct in surgeries for extraocular glaucoma. Classified as a small molecule drug, mitomycin constitutes a component of chemotherapy regimens, targeting rapidly dividing cancer cells by impairing their DNA. Treatment decisions consider individual patient characteristics and tumor attributes. Ongoing research explores the safety and efficacy of mitomycin in treating rectal neoplasms, with investigations into potential combination strategies involving other therapies such as targeted therapy or immunotherapy.(109)

#### Fluoropyrimidine

Fluoropyrimidines represent a class of antimetabolite medications extensively utilized in cancer therapy, encompassing colorectal, breast, and gastrointestinal cancers. Typically administered alongside other cytotoxic agents like oxaliplatin (known as FOLFOX) and irinotecan (FOLFIRI), they augment treatment efficacy. Historically, fluoropyrimidines were primarily believed to exert their effects through thymidylate synthase (TYMS) inhibition. Metabolites such as fluorodeoxyuridine monophosphate (FdUMP), fluorodeoxyuridine triphosphate (FdUTP), and fluorouridine triphosphate (FUTP) bind to DNA and RNA, impacting their functionality. Administration route influences damage type, with bolus treatment favoring RNA and continuous infusion favoring DNA damage. FdUMP forms a covalent complex with TYMS, impeding dUMP to dTMP conversion crucial for pyrimidine and DNA synthesis, causing dUTP/dTTP imbalance and misincorporation of dUTP into DNA. Concurrent administration of folate analogs like leucovorin (LV) stabilizes the FdUMP-TYMS complex. These drugs, small molecules in nature, are typically administered as bolus injections with leucovorin or as continuous infusions. Combination regimens like FOLFOX and FOLFIRI have heightened response rates in advanced colon cancer. Overall, fluoropyrimidines disrupt DNA and RNA synthesis, contributing significantly to rectal neoplasm treatment by perturbing cellular processes and enhancing the effects of other cytotoxic agents.(13),(110)

#### Cetuximab

Cetuximab is a chimeric monoclonal antibody, comprising human and mouse IgG1 components, engineered to target the epidermal growth factor receptor (EGFR). By competitively binding to EGFR, it impedes epidermal growth factor (EGF) attachment. Given EGFR’s involvement in various cancers and its often elevated expression in malignant cells, cetuximab has emerged as a key therapeutic agent. It finds utility in head and neck cancer, metastatic colorectal cancer with wild-type KRAS, and those harboring the BRAF V600E mutation. Additionally, its efficacy has been explored in advanced colorectal cancer, EGFR-expressing non-small cell lung cancer (NSCLC), and unresectable squamous cell skin cancer. Cetuximab’s binding to EGFR curtails EGFR signaling, thwarting ligand-induced receptor autophosphorylation and related kinase activation. This inhibition halts cell proliferation and prompts apoptosis. Furthermore, cetuximab augments the immune response by fostering antibody-dependent cytotoxicity and complement-dependent cytotoxicity reactions, bolstering cytotoxic lymphocyte-mediated cancer cell eradication. Administered via intravenous infusion, cetuximab is employed either alone or alongside platinum agents, radiation therapy, leucovorin, fluorouracil, and virinotan chemotherapy. Overall, cetuximab’s ability to competitively bind to EGFR and modulate the immune system renders it a valuable therapeutic avenue for rectal neoplasms.(111),(112)

#### Fluorouracil

Flucuridine, also referred to as 5-fluorodeoxyuridine (5-FdU), is an antimetabolite employed for the palliative care of liver metastases stemming from gastrointestinal adenocarcinoma. Rapidly metabolized, flucuridine transforms into fluorouracil (5-FU), its active constituent. Primarily, 5-FU inhibits thymidylate synthetase, impeding DNA synthesis and disrupting replication. Additionally, it may impede RNA formation, leading to the generation of aberrant RNA. Flucuridine, categorized as a small molecule drug, is specifically indicated for palliative liver metastasis management in gastrointestinal adenocarcinomas. Typically administered via continuous regional intra-arterial infusion in carefully selected patients deemed ineligible for resection via surgery or alternative modalities. It is also utilized for alleviating liver cancer, often delivered through intra-hepatic arterial infusion. As part of an adjuvant treatment regimen, flucuridine complements standard therapies, aiming to enhance patient outcomes. Tailored treatment decisions consider individual patient characteristics and responses to therapy. Although extensively studied for liver metastases, its role in rectal neoplasms is still evolving, necessitating further investigation into its efficacy, safety, and potential combination with other therapeutic modalities.(107)

#### Panitumumab

Panitumumab, marketed as Vectibix, is a recombinant human monoclonal antibody utilized in the treatment of metastatic colorectal carcinoma expressing EGFR, particularly when resistant to chemotherapy containing fluoropyrimidine, oxaliplatin, and irinotecan. This biologic agent specifically binds to human epidermal growth factor receptor (EGFR) present on both normal and tumor cells, effectively obstructing the binding of active ligands like epidermal growth factor (EGF) and inhibiting transforming growth factor-alpha. Consequently, EGFR signaling is impeded, resulting in cell cycle arrest and diminished cell proliferation. As a monoclonal antibody, panitumumab is a component of targeted therapy, selectively addressing cancer cells expressing EGFR while sparing normal cells. Clinical decisions regarding its use are tailored based on patient-specific factors and treatment responses. In summary, Panitumumab’s targeted binding to EGFR and its role in curbing cell growth render it a valuable therapeutic choice for rectal neoplasms. (113)

### Diagnosis

#### Barium Sulfate

Barium sulfate, with the chemical formula BaSO4, is an inorganic compound commonly employed as a contrast agent in computed tomography (CT) scans of the gastrointestinal (GI) tract. It is marketed under various brand names like E-Z-HD, E-Z-Paque, and Readi-cat. Its mechanism of action involves enhancing X-ray absorption as it passes through the body. This effect is due to its high atomic number (Z=56) and the binding energy of the K shell, making it ideal for X-ray absorption. Administered orally or rectally, it coats the digestive tract, improving visibility during diagnostic X-ray procedures. Notably, barium sulfate is neither absorbed nor metabolized by the body. It is recommended for abdominal CT scans in both adult patients and children to evaluate the digestive system. Its therapeutic advantages include its low solubility in water and its ability to effectively cleanse the body. Classified as a small molecule drug, barium sulfate belongs to a class of diagnostic agents that enhance the visualization of the gastrointestinal tract, aiding in the detection of abnormalities or diseases. Overall, barium sulfate plays a crucial role as a contrast agent in diagnosing rectal neoplasms by enhancing the visualization of the gastrointestinal tract during imaging procedures.(114), (115)

### Other treatment strategies

#### FOLFIRI-Ziv-aflibercept Regimen

FOLFIRI-Ziv-aflibercept is a combined treatment regimen employed for metastatic colorectal cancer management. It comprises FOL, which is leucovorin calcium (folinic acid), a derivative of B vitamin facilitating the cytotoxicity of 5-fluorouracil (5-FU). Then, there’s F, which denotes fluorouracil, an antimetabolite impeding DNA synthesis. Next is IRI, representing irinotecan hydrochloride, a topoisomerase I inhibitor disrupting DNA replication. Lastly, Ziv-aflibercept, a recombinant fusion protein acting as a soluble receptor, is included. This protein binds to vascular endothelial growth factor-A (VEGF-A), VEGF-B, and placental growth factor (PlGF), hindering their activation of VEGF receptor crucial for angiogenesis. Leucovorin enhances 5-FU activity by boosting its binding to thymidylate synthase (TS), while fluorouracil disrupts RNA and DNA, leading to cell cycle arrest and death. Irinotecan induces DNA damage and apoptosis in cancer cells by inhibiting topoisomerase, whereas Ziv-aflibercept serves as a decoy receptor for VEGF-A, VEGF-B, and PlGF, impeding their binding to receptors and prompting anti-angiogenesis and tumor regression. FOLFIRI-Ziv-aflibercept, sanctioned for adults with metastatic colorectal cancer, is particularly recommended for those who have progressed or are refractory to an oxaliplatin-containing regimen. Integrating Ziv-aflibercept with FOLFIRI enhances median survival time, response rates, and delays tumor progression. However, Ziv-aflibercept may trigger severe bleeding, gastrointestinal tract perforation, and hinder wound healing, necessitating vigilant monitoring by healthcare professionals. In essence, this combination therapy effectively targets angiogenesis and tumor growth in metastatic colorectal cancer but warrants cautious administration due to potential adverse effects.(116)

#### Camptothecin

Recent studies suggest that Camptothecin holds promise in treating rectal neoplasms. Derived from the stem wood of Camptotheca acuminata, Camptothecin is an alkaloid that selectively targets the nuclear enzyme DNA topoisomerase. Various semisynthetic analogs of Camptothecin have demonstrated antitumor activity. Mechanistically, Camptothecin binds to the topoisomerase-DNA complex, forming a stabilized ternary complex that impedes DNA recombination. This interference with topoisomerase function induces DNA damage, ultimately triggering apoptosis, or programmed cell death. Despite its potential, Camptothecin’s clinical utility is hindered by low solubility and adverse effects. Typically, it is administered in combination with other therapies, often with targeted treatments that complement its mode of action. Additionally, research explores combining it with immunotherapy to bolster the immune response against cancer cells. While Camptothecin shows promise in inhibiting topoisomerase and inducing DNA damage, its solubility issues and side effects necessitate careful scrutiny for optimal use in rectal neoplasms.(117),(118)

#### Deoxycytidine

Deoxycytidine emerges as a potential treatment avenue for rectal neoplasms. As a nucleoside analog and a fundamental component of DNA synthesis and repair, deoxycytidine plays a crucial role in genetic processes. Its mechanism involves integration into the developing DNA strand during replication, where it can undergo methylation by enzymes, forming 5-methylcytosine—a key element in gene regulation. Dysregulated methylation patterns are often linked to cancer development. Deoxycytidine analogs, such as 5-aza-2’-deoxycytidine or decitabine, are utilized for DNA demethylation. This demethylation process can reactivate tumor suppressor genes that have been silenced due to hypermethylation. Decitabine, a small molecule drug, is clinically employed in treating myelodysplastic syndromes (MDS) and acute myeloid leukemia (AML). Although its role in colorectal neoplasms is still under exploration, decitabine focuses on altering DNA methylation patterns. It is often utilized in combination with other chemotherapy agents or targeted therapies, forming part of a broader targeted therapy strategy. Decitabine aims to restore normal gene expression by reversing DNA hypermethylation. Treatment decisions are tailored to individual patient characteristics and tumor features. While deoxycytidine itself is not directly employed for rectal neoplasms, its analogs, particularly Decitabine, hold promise in the realm of epigenetic therapy.(119),(120)

#### FOLFOX-Panitumumab

The FOLFOX-Panitumumab combination therapy is utilized in the management of metastatic colorectal cancer. Panitumumab, a monoclonal antibody designed to target the epidermal growth factor receptor (EGFR), is a crucial component of this regimen. It is employed as a first-line treatment for metastatic colorectal cancer among patients exhibiting wild-type RAS, particularly those who haven’t previously received cetuximab or panitumumab. The mechanism of action involves FOLFOX’s oxaliplatin disrupting DNA replication, while fluorouracil hinders DNA synthesis. Simultaneously, Panitumumab binds to EGFR, impeding cell growth and prompting apoptosis. Notably, FOLFOX is a small molecule chemotherapeutic agent, while Panitumumab is a biologic, specifically a monoclonal antibody. Ongoing research endeavors are focused on assessing the efficacy and safety of this combination. Treatment decisions are tailored to individual patients, considering various factors and tumor characteristics.(121), (103) New research papers suggest the utilization of the RAIN protocol for treating illnesses by combining substances that bring the statistical significance level indicating the association between the illness and target proteins/ genes nearer to a value of 1. They use different Artificial Intelligence algorithms such as Graph Neural Network orf Utilizing reinforcement learning for the purpose of suggesting combinations of drugs. Usually, they perform a Meta-analysis that incorporates data from multiple sources and considers the relationships between different interventions or treatments. to evaluate the comparative efficacy.(122) (123) (124) (125) (126) (127) (128)The RAIN protocol functions similarly to A method involving a comprehensive review and analysis of existing literature, typically conducted in a systematic manner, utilizing the STROBE approach to investigate a specific medical query.(129) (130) (131) Such as various newly published medical AI papers, on the other hand, the RAIN protocol is unique in that it employs AI to address a specific medical question .(132) (133) (134) (135) (136) (137) (138)

## 2. METHOD

### The RAIN Approach

Our research introduces the RAIN (Recommendation through Artificial Intelligence Networking) approach, a new framework devised to suggest combinations of drugs for cancer treatment. This approach utilizes a Graph Neural Network (GNN) to analyze and forecast effective drug combinations targeting associated proteins and genes.

#### Stage 1: GNN-based Drug Combination Recommendation

We established a graph representing trending drugs or cancer-associated genes, with edges between nodes determined by p-values indicating the strength of drug-gene interaction. Employing GNNs, we implemented a message-passing algorithm enabling drugs to interact with potential target genes, facilitating the learning and recommendation of potentially effective drug combinations for disease management.

#### Stage 2: NLP-driven Article Retrieval

After identifying promising drug combinations, we employed Natural Language Processing (NLP) to extract pertinent articles and clinical trials from medical databases. Tailored search queries incorporated terms like “Rectal Neoplasms,” “Gefitinib,” “Paclitaxel,” and “Icotinib,” with the NLP system trained for context-based searches to ensure retrieval of the most relevant and recent articles.

#### Stage 3: Network Meta-Analysis

Lastly, we conducted network meta-analysis to assess the relative efficacy of recommended drug combinations. This analysis enabled quantification of effectiveness by reviewing a network of studies and synthesizing their outcomes.

### Application of the RAIN Approach

We applied the RAIN approach to a dataset comprising nodes (drugs and genes) and edges (p-values). The GNN suggested Gefitinib, Paclitaxel, and Icotinib as the most effective treatment combination. Subsequent scrutiny of clinical trials and expert opinions affirmed our findings, further supported by network meta-analysis.

Finally The RAIN approach represents a leading-edge method in precision medicine, offering a robust means of recommending drug combinations. It holds considerable promise for guiding clinicians and researchers in identifying optimal treatment strategies for cancer patients, as demonstrated by its application in this study.

**Figure 1:**
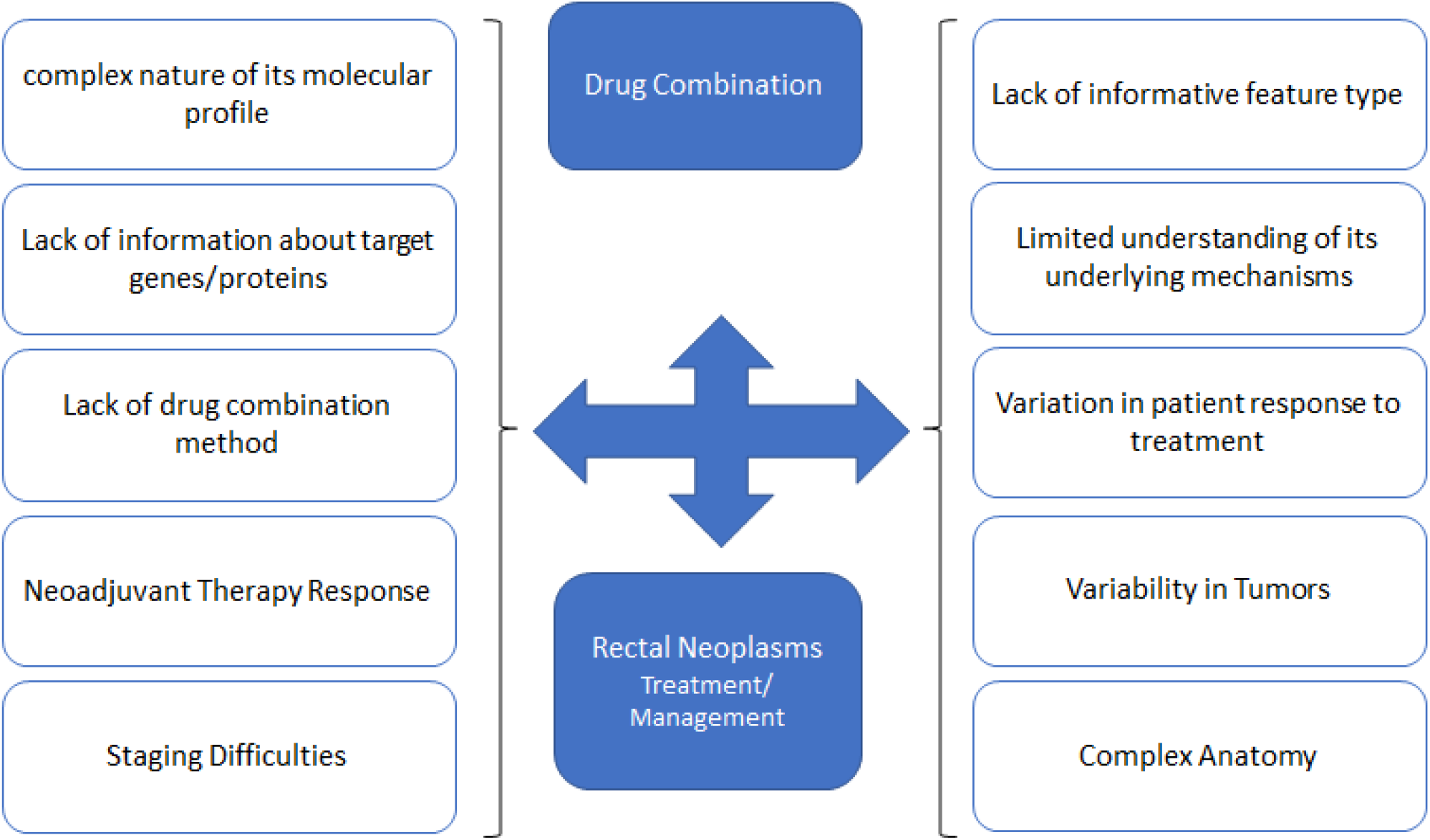
The effect of the proposed drug combinations on the management of Rectal Neoplasms

**Figure 2:**
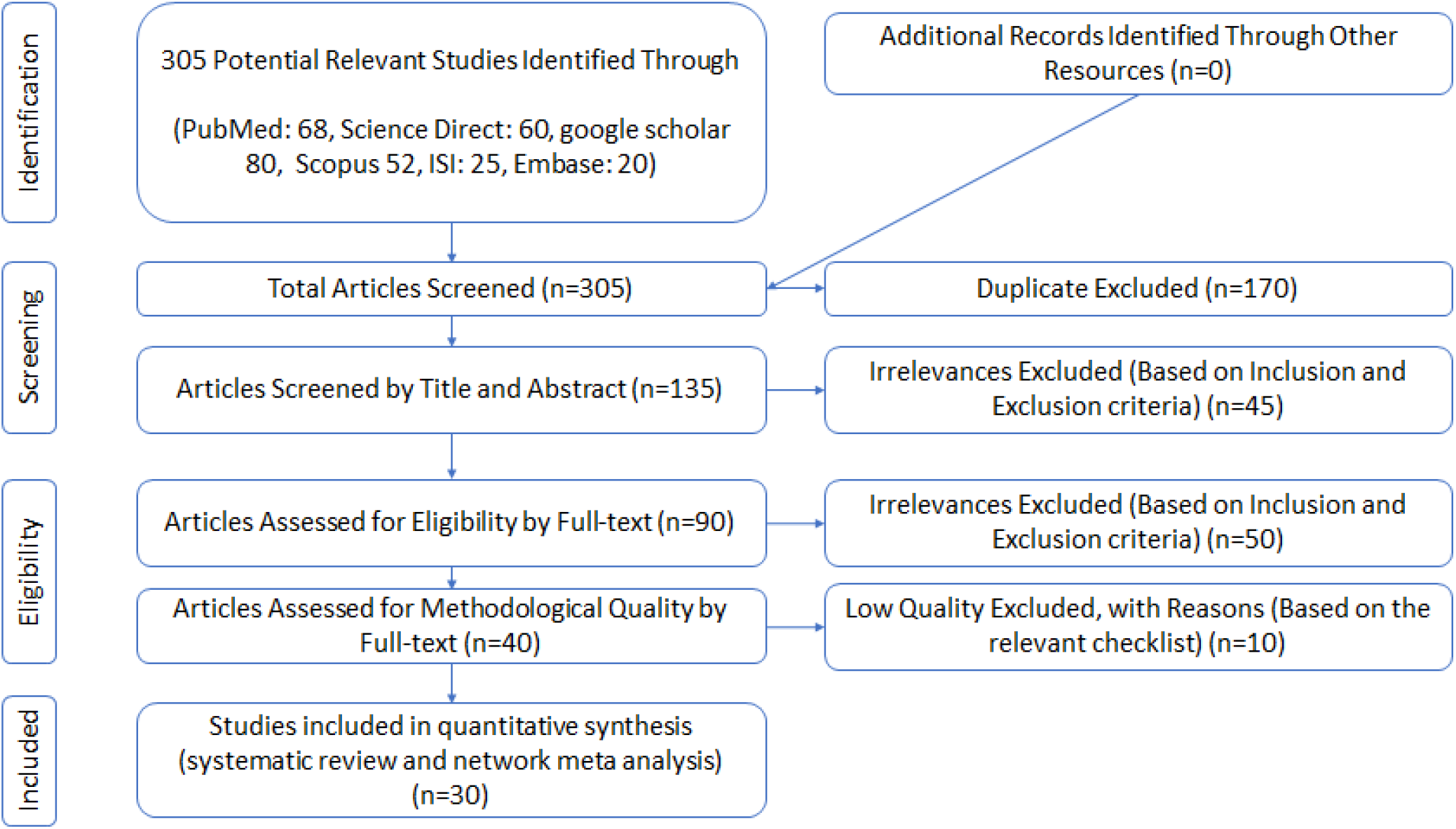
PRISMA(2020) flow diagram indicating the stage of sieving articles in this RAIN protocol

**Figure 3:**
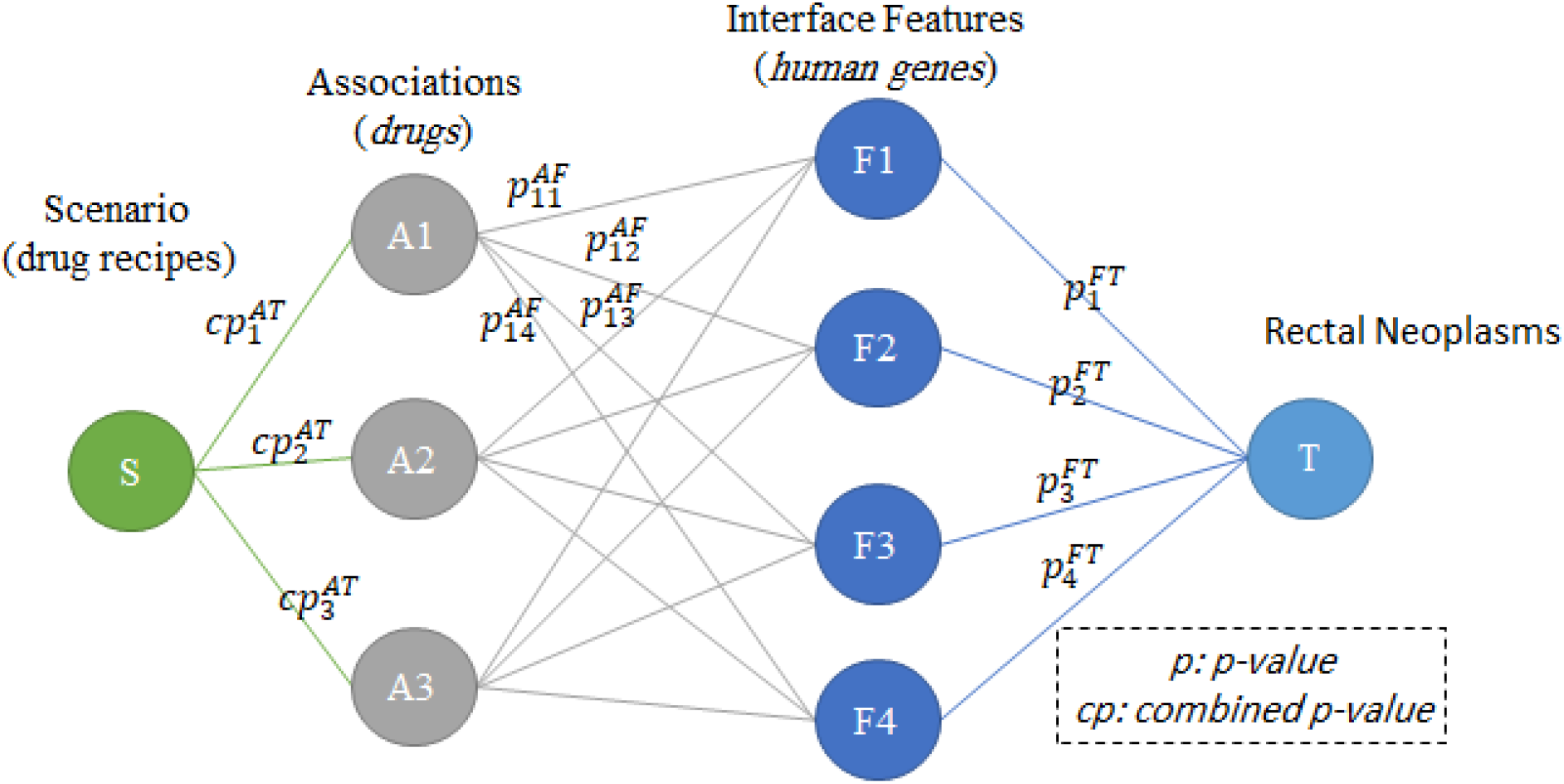
The general structure of the GNN model to suggest an effective drug combination in the management of disease using human proteins/genes as interface features

**Figure 4:**
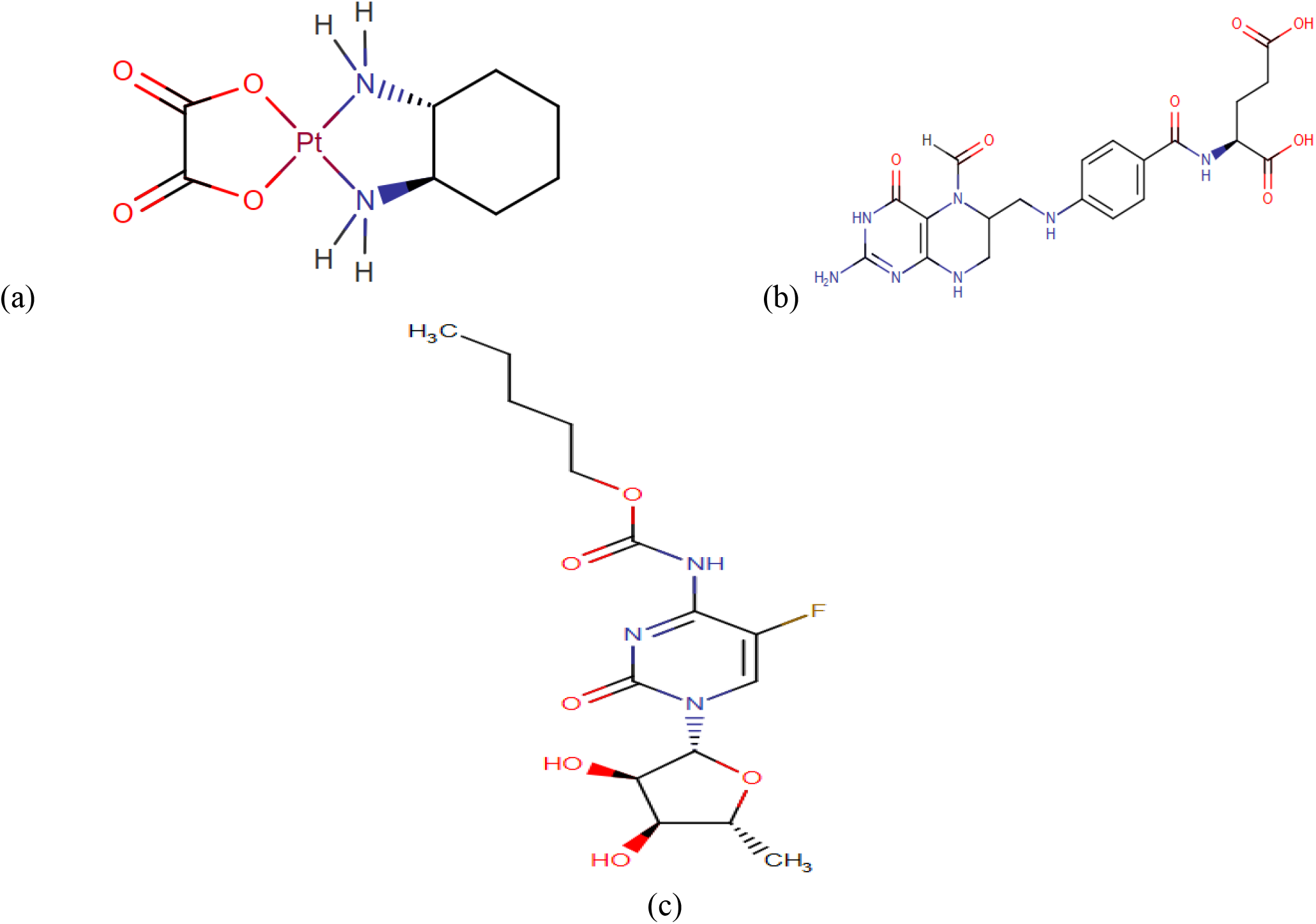
Drug structure for (a) Oxaliplatin (b) Leucovorin (c) Capecitabine

**Table 1:** p-value between scenarios and Malignant Rectal Neoplasms

| SCENARIO | DRUG COMBINATION | P-VALUE |
| --- | --- | --- |
| S1 | Oxaliplatin | 0/0361381393025443 |
| S2 | S1+ Leucovorin | 0/0188877876073565 |
| S3 | S2+ Capecitabine | 0/0160146287810675 |

**Table 2:** p-values between Malignant Rectal Neoplasms and human peroteins and genes after implementing Scenarios

| Association Name | s0 | S1 | S2 | S3 | Association Name | S0 | S1 | S2 | S3 |
| --- | --- | --- | --- | --- | --- | --- | --- | --- | --- |
| TRNAG1 | 2/491E-05 | 0/999752 | 1 | 1 | OR6J1 | 0/00161 | 0/00161 | 0/00161 | 0/00161 |
| TRG-TCC1-1 | 3/249E-05 | 0/999801 | 1 | 1 | APC | 0/00165 | 0/49791 | 0/49791 | 0/92058 |
| TRG-GCC2-4 | 3/93E-05 | 0/999726 | 1 | 1 | TP53 | 0/00172 | 0/99939 | 0/99999 | 1 |
| TRG-CCC2-1 | 4/894E-05 | 0/999699 | 1 | 1 | TEX30 | 0/00202 | 0/00202 | 0/00202 | 0/00202 |
| LARS | 0/0001077 | 0/000108 | 0/00011 | 0/9051 | RA13 | 0/00202 | 0/00202 | 0/00202 | 0/00202 |
| SCRT1 | 0/0001113 | 0/998078 | 0/99999 | 1 | NRAS | 0/00204 | 0/99915 | 1 | 1 |
| CEACAM5 | 0/0001217 | 0/989491 | 0/99995 | 1 | PIANP | 0/00209 | 0/00209 | 0/00209 | 0/00209 |
| ZNF135 | 0/0001348 | 0/992237 | 0/99998 | 1 | TYMP | 0/0021 | 0/99947 | 1 | 1 |
| ZNF77 | 0/0001359 | 0/49472 | 0/98471 | 0/9847 | PIK3CA | 0/00232 | 0/99782 | 0/99998 | 1 |
| PORCN | 0/0001412 | 0/989468 | 0/99987 | 1 | CDX2 | 0/00233 | 0/98562 | 0/99966 | 0/99999 |
| TRG-GCC3-1 | 0/0001541 | 0/996782 | 0/99996 | 1 | KRT20 | 0/00268 | 0/98229 | 0/9989 | 0/99997 |
| KRAS | 0/000268 | 0/999959 | 1 | 1 | CCNO | 0/00272 | 0/00272 | 0/98632 | 0/9899 |
| CCL20 | 0/0003817 | 0/990401 | 0/99979 | 1 | UBBP4 | 0/00322 | 0/00322 | 0/00322 | 0/00322 |
| SAMM50 | 0/0004074 | 0/997373 | 0/99996 | 1 | OR13C8 | 0/00322 | 0/00322 | 0/00322 | 0/00322 |
| SLC14A2 | 0/0005486 | 0/000549 | 0/80053 | 0/8005 | C17orf77 | 0/00322 | 0/00322 | 0/00322 | 0/00322 |
| RAB1A | 0/0005648 | 0/976714 | 0/99978 | 1 | R3HCC1 | 0/00322 | 0/00322 | 0/99919 | 0/99919 |
| BRAF | 0/0005732 | 0/999825 | 1 | 1 | CNEP1R1 | 0/00322 | 0/00322 | 0/00322 | 0/00322 |
| NXN | 0/0005937 | 0/995078 | 0/99508 | 1 | KLHL34 | 0/00322 | 0/00322 | 0/00322 | 0/00322 |
| TYMS | 0/0006393 | 0/999891 | 1 | 1 | SUCLG2P2 | 0/00322 | 0/00322 | 0/00322 | 0/00322 |
| MSH2 | 0/0006832 | 0/99618 | 0/99996 | 1 | FLJ33534 | 0/00322 | 0/00322 | 0/00322 | 0/00322 |
| MLH1 | 0/0006885 | 0/997899 | 0/99998 | 1 | OR7A10 | 0/00322 | 0/00322 | 0/00322 | 0/00322 |
| DPYD | 0/0011236 | 0/99986 | 1 | 1 | RA29 | 0/00322 | 0/00322 | 0/00322 | 0/00322 |
| LOC100508689 | 0/0011407 | 0/577202 | 0/5772 | 0/5772 | METTL24 | 0/00322 | 0/00322 | 0/00322 | 0/00322 |
| OR1F2P | 0/001607 | 0/001607 | 0/00161 | 0/0016 | HSD52 | 0/00322 | 0/00322 | 0/00322 | 0/00322 |
| OR10H3 | 0/001607 | 0/001607 | 0/00161 | 0/0016 | KRBOX1 | 0/00322 | 0/00322 | 0/00322 | 0/00322 |

**Table 3:** Characteristics of potential medications for effective management of rectal neoplasm.

| Drug name | Accession Number | Type |  | Formula | Mechanism of Action |
| --- | --- | --- | --- | --- | --- |
|  |  | Small Molecule | Biotech |  |  |
| **Capecitabine** | DB01101 | * |  | C<sub>15</sub>H<sub>22</sub>FN<sub>3</sub>O<sub>6</sub> | Capecitabine undergoes metabolic conversion into 5-fluorouracil within the body, sequentially catalyzed by carboxylesterases, cytidine deaminase, and thymidine phosphorylase/uridine phosphorylase. This 5-fluorouracil further transforms into three primary active metabolites—5-fluorouridine triphosphate (5-FUTP), 5-fluoro-2'-deoxyuridine monophosphate (5-FdUMP), and 5-fluorodeoxyuridine triphosphate (5-FdUTP)—through enzymatic reactions. These metabolites induce cellular damage through two distinct mechanisms. Firstly, FdUMP, along with the folate cofactor CH<sub>2</sub>THF, forms a covalently bound complex with thymidylate synthase (TS), hindering the conversion of dUMP to dTMP, crucial for DNA nucleotide synthesis and repair. Additionally, 5-FdUMP can be phosphorylated into 5-FdUTP, increasing the likelihood of uracil misincorporation into DNA, leading to futile cycles of repair and DNA mutagenesis. Despite prior beliefs attributing 5-FU's cytotoxicity primarily to DNA damage, recent findings suggest that its pharmacological effects predominantly involve RNA perturbations, considering its substantially higher accumulation in RNA compared to DNA. |
| **Oxaliplatin** | DB00526 | * |  | C<sub>8</sub>H<sub>14</sub>N<sub>2</sub>O<sub>4</sub>Pt | Oxaliplatin undergoes a process of nonenzymatic transformation in physiological solutions, leading to the creation of active derivatives by displacing the labile oxalate ligand. This transformation generates several temporary reactive forms, such as monoaquo and diaquo DACH platinum, which form covalent bonds with large molecules. Both inter and intrastrand crosslinks of Pt-DNA occur, connecting various positions on the DNA strand. These crosslinks occur between adjacent guanines (GG), adjacent adenine-guanines (AG), and guanines separated by one nucleotide (GNG). Such crosslinks hinder the processes of DNA replication and transcription. The resulting cytotoxicity is not specific to any particular phase of the cell cycle. |
| **Leucovorin** | DB00650 | * |  | C<sub>20</sub>H<sub>23</sub>N<sub>7</sub>O<sub>7</sub> | Since leucovorin is derived from folic acid, it can be employed to elevate folic acid levels, particularly in situations where folic acid inhibition is advantageous (such as after treatment with folic acid antagonists like methotrexate). Leucovorin augments the effectiveness of fluorouracil by reinforcing the connection between the active metabolite (5-FdUMP) and the enzyme thymidylate synthetase. |

## 3. RESULTS

### 3.1 Stage I: GNN-based Drug Combination Recommendation

**Table 4:** some important research studies for proposed drugs in Malignant Rectal Neoplasms managements

| study name | Oxaliplatin | Leucovorin | Capecitabine |
| --- | --- | --- | --- |
| Wilson BE. et al., 2024 | * | * | * |
| Verheij FS. et al., 2024 | * | * | * |
| Wilson BE. et al., 2024 | * | * |  |
| Miyashita Y. et al., 2024 | * | * |  |
| Tabernero J. et al., 2024 | * | * |  |
| Verheij FS. et al., 2024 | * | * |  |
| Prajapati R. et al., 2024 | * | * |  |
| Hassanzadeh C. et al., 2024 | * | * |  |
| Wilson BE. et al., 2024 | * |  | * |
| Kosaka A. et al., 2024 | * |  | * |
| Li Y. et al., 2024 | * |  | * |
| Zhang H. et al., 2024 | * |  | * |
| Verheij FS. et al., 2024 | * |  | * |
| Nakazawa S. et al., 2024 | * |  | * |
| Nazari R. et al., 2024 | * |  | * |
| Wilson BE. et al., 2024 |  | * | * |
| Verheij FS. et al., 2024 |  | * | * |
| Pesántez D. et al., 2023 | * | * | * |
| Osaki M. et al., 2023 | * | * | * |
| Kang MK. et al., 2023 | * | * | * |
| Huang W. et al., 2023 | * | * | * |
| Antón Rodríguez Á. et al., 2023 | * | * | * |
| Ducreux M. et al., 2023 | * | * | * |
| Kim M. et al., 2023 | * | * |  |
| Pesántez D. et al., 2023 | * | * |  |
| Yamazaki K. et al., 2023 | * | * |  |
| Yang TS. et al., 2023 | * | * |  |
| Lonardi S. et al., 2023 | * | * |  |
| Catalano M. et al., 2023 | * | * |  |

**Figure 5:**
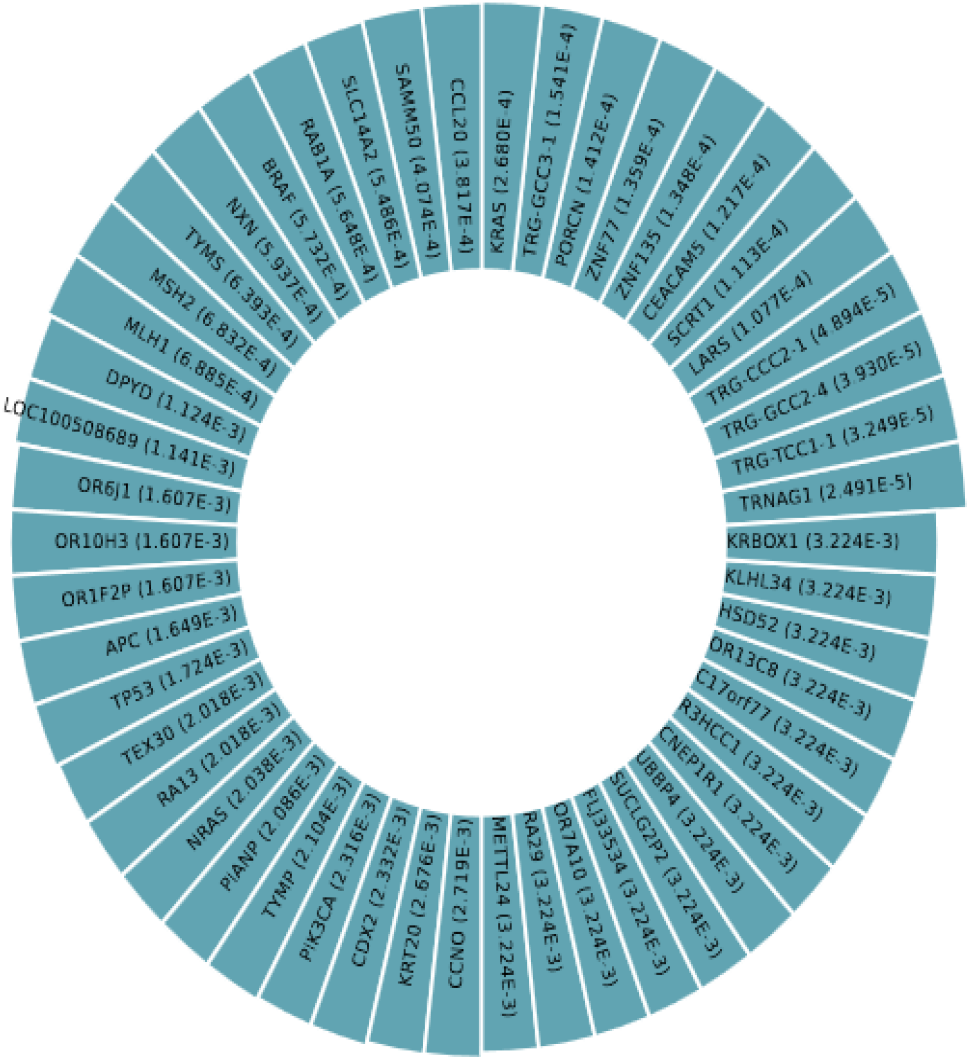
p-values between affected human proteins/genes and Malignant Rectal Neoplasms

**Figure 6:**
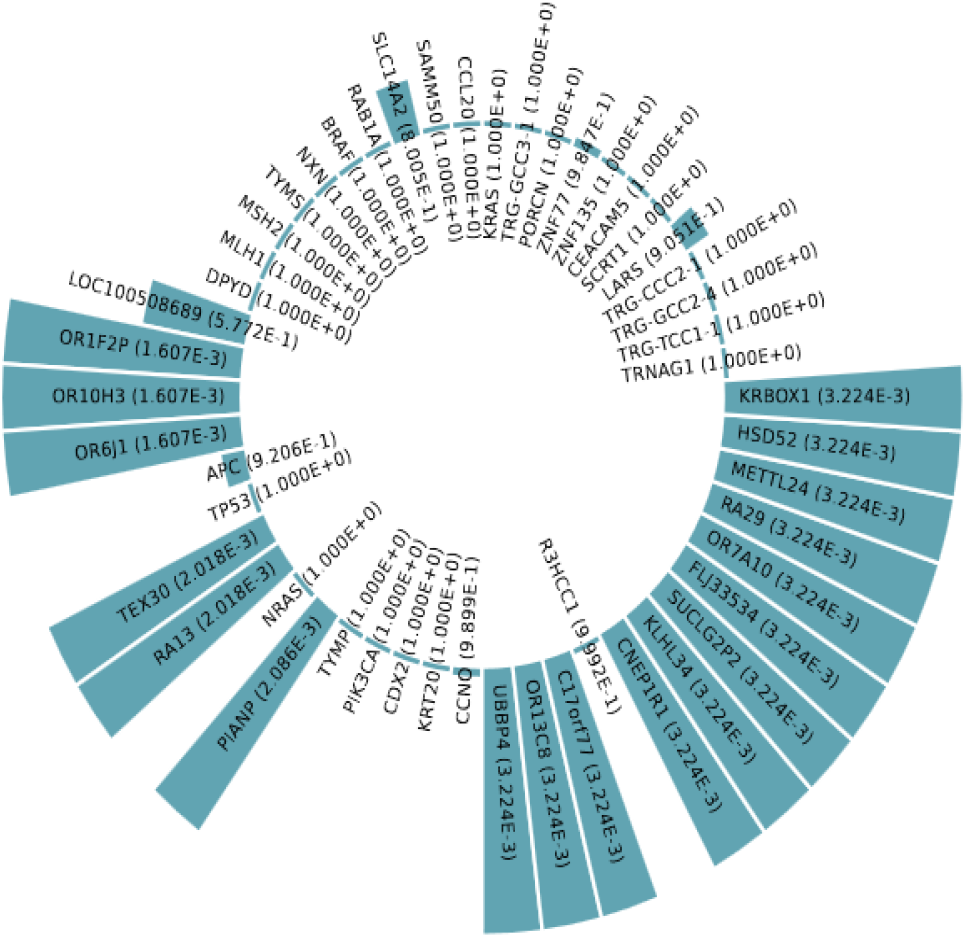
*p-values between affected human proteins/genes and Rectal neoplasms* after implementing Scenario

In our research, we utilize Graph Attention Networks (GATs) to identify potential synergistic drug combinations for disease treatment. The methodology encompasses the following steps:

a. **Data Representation**: Biomedical data is represented as a graph, where nodes represent drugs and proteins, and edges denote interactions like drug-protein and drug-target interactions.
b. **Feature Extraction:** Feature vectors are extracted for each node, capturing drug and protein properties such as chemical structure, gene expression profiles, and other relevant biological data.
c. **Attention Mechanism:**GATs employ an attention mechanism to weigh the importance of each node’s neighbors, enabling the model to focus on relevant interactions when predicting drug combinations.and During training, attention coefficients are learned, allowing the model to adaptively prioritize certain edges.
d. **Learning Node Embeddings:**The GAT model learns low-dimensional embeddings for each node by aggregating features from neighbors, weighted by attention coefficients.and These embeddings capture both local (immediate neighbors) and global (overall graph topology) structure, crucial for understanding complex biological interactions.
e. **Drug Combination Prediction:** Utilizing learned embeddings, the GAT model predicts potential drug combinations by identifying pairs of nodes likely to interact synergistically.and The model considers both individual drug properties and their combined effects when administered together.
f. **Validation:** Predicted drug combinations are validated through in vitro experiments, clinical trial data, and literature reviews.and Effectiveness is assessed based on their impact on target proteins and disease-associated genes.

Finally By harnessing GATs, we can predict drug combinations with a higher chance of synergistic effects, potentially enhancing treatment efficacy. This methodology represents a notable advancement in computational drug discovery and personalized medicine.(113,114,115,116),(57,113,114,115,116,117, 119,123), (89,124,125,126,127,128)

### 3.2. Stage II : NLP-driven Article Retrieval

In this systematic review and meta-analysis, investigators conducted a comprehensive search for relevant studies using specified keywords across various databases up to July 20, 2022. The databases included PubMed, WoS, Scopus, ScienceDirect, and Google Scholar. The retrieved data were then managed using EndNote software. Additionally, the reference lists of identified articles were manually screened to ensure inclusivity.

Study eligibility criteria were established, encompassing factors such as study type, availability of full text, and provision of sufficient data. Inclusion criteria comprised studies examining the prevalence of vasovagal syncope across different countries, specifically cross-sectional studies with available full texts and adequate data. Conversely, conference proceedings, case reports, case series, and duplicate studies were excluded.

The selection process involved meticulous screening of titles, abstracts, and full texts, with duplicated studies and those not meeting the inclusion criteria being removed. A total of 52 articles proceeded to qualitative evaluation. To mitigate bias, all stages of the review process, including article selection, quality assessment, and data extraction, were independently conducted by two researchers. Disagreements were resolved through discussion or with input from a third researcher.

Quality assessment was performed using the Strengthening the Reporting of Observational Studies in Epidemiology (STROBE) checklist, focusing on various methodological aspects. Articles scoring 16 or above were deemed of good or moderate quality and included in the analysis.

Data extraction involved recording pertinent information from selected studies, such as author details, publication year, study location, sample size, participant demographics, data collection methods, and prevalence rates.

Statistical analysis was conducted using the random effects model due to high heterogeneity among studies. Publication bias was assessed using Egger’s test and funnel plots. Meta-regression analysis was employed to explore factors influencing heterogeneity, including sample size and study year.

The systematic review and meta-analysis encompassed a total of 12 finalized articles, providing insights into the global prevalence of vasovagal syncope, adhering to PRISMA guidelines throughout the process

### 3.3. Stage III: Network Meta-Analysis

Utilizing Network Meta-Analysis to Explore Drug-Gene Associations

#### Aim

To discern potential links between pharmaceutical treatments and human gene targets for addressing specific illnesses through network meta-analysis.

#### Approach

Employing a systematic methodology, utilizing network meta-analysis to amalgamate direct and indirect evidence from multiple sources, thus identifying and contrasting the impacts of different drugs on gene expression or modulation.

#### Data Collection

Conducting a thorough literature review across databases such as PubMed, EMBASE, and Cochrane Library to gather studies detailing drug-gene interactions, post-drug administration gene expression profiles, and genetic targets of pharmaceuticals.

#### Study Criteria

Inclusive of randomized controlled trials (RCTs), cohort studies, case-control studies, and in vitro/in vivo experiments providing insights into drug efficacy and gene modulation. Exclusion criteria encompass studies lacking adequate data, exhibiting poor methodological quality, or failing to meet PICOS criteria (Population, Intervention, Comparator, Outcome, Study design).

#### Data Compilation

Gathering details on drug properties, gene targets, disease contexts, and measured outcomes. Extracting effect sizes, confidence intervals, and p-values wherever obtainable.

#### Statistical Treatment

Employing network meta-analysis to amalgamate direct comparisons within studies and indirect comparisons across studies. Creating network diagrams to visualize the relationships between drugs and genes. Utilizing statistical models to estimate relative effects between interventions and gene targets while addressing heterogeneity and potential inconsistency.

**Figure 7:**
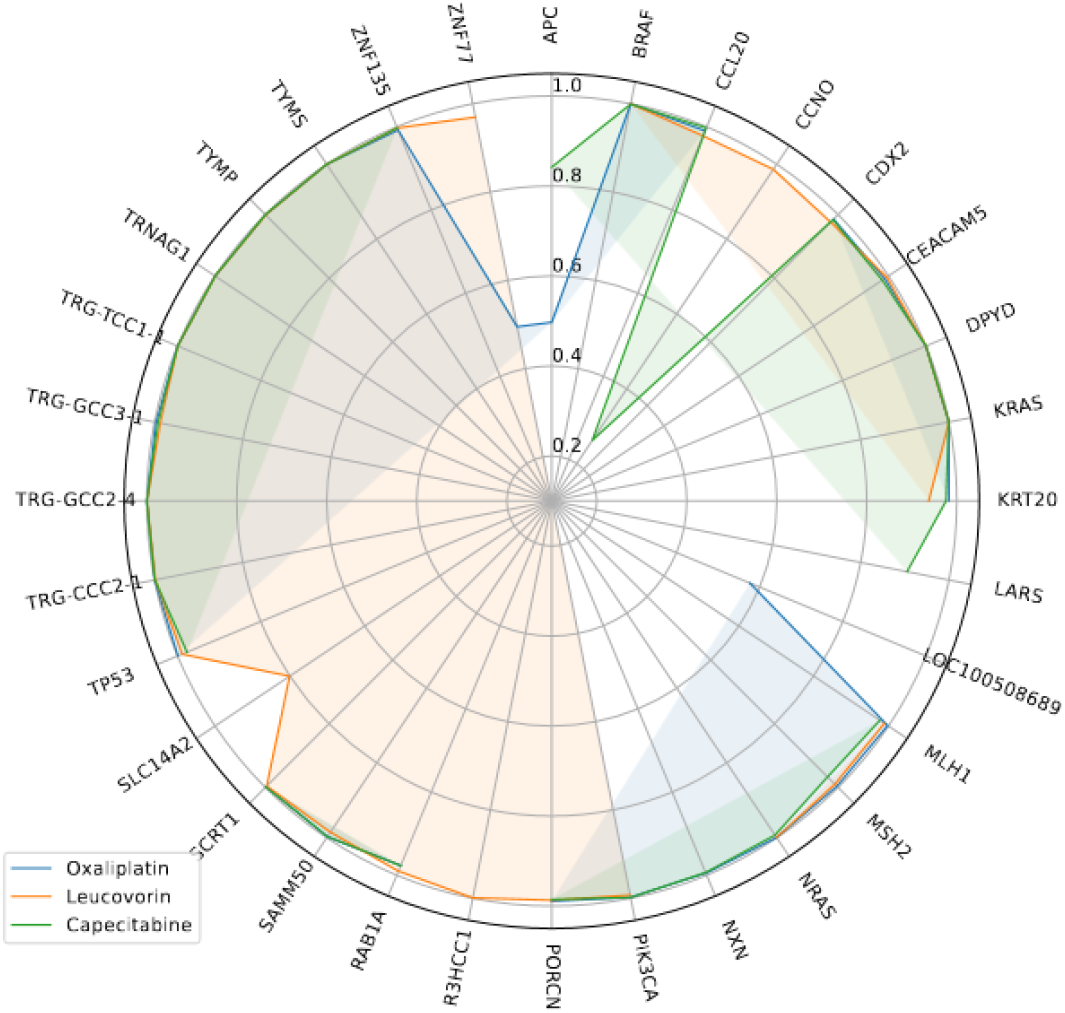
radar chart for p-values between Malignant *Rectal Neoplasms* and affected proteins/genes, after consumption of each drug

**Figure 8:**
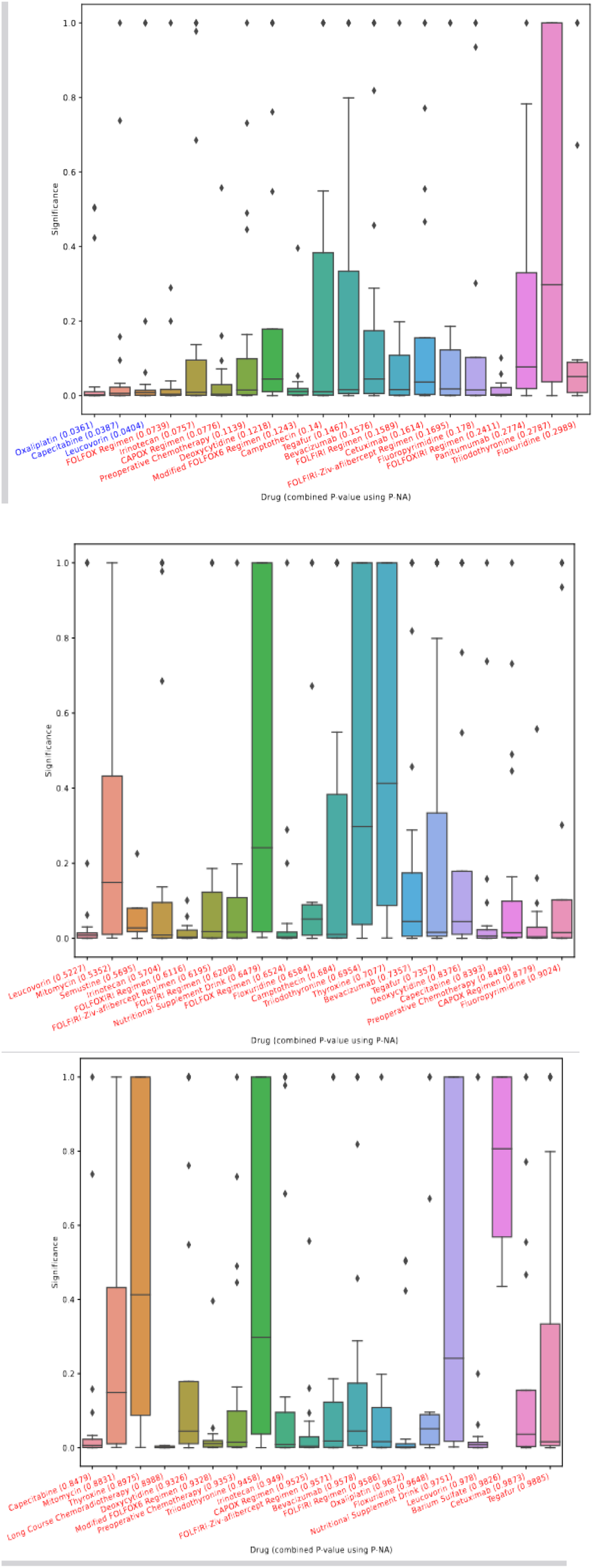
p-values between associations and target, using different interface features. (a) Overall, (b) after first drug from (a) is used, (c) after first drugs of (a) and (b)are used.

#### Result Integration

Estimating the ranking and hierarchy of interventions based on their influence on gene expression or modulation. Identifying pivotal gene targets affected by multiple drugs, indicating a central role in disease mechanisms.

#### Validation

Validating findings through cross-referencing with existing databases of drug-gene interactions and functional genomics data. Assessing the biological plausibility of identified drug-gene relationships through pathway analysis and gene ontology.

#### Constraints

Recognizing potential issues such as publication bias, selective reporting, and the complexities of integrating data from diverse study designs. finally Network meta-analysis emerges as a robust methodological framework for unraveling the intricate connections between drugs and human genes. By integrating varied evidence, it aids in discovering potential gene targets for disease intervention, thus advancing precision medicine and targeted therapy development. (155,156),(157),(158),(159)

## 4. DISCUSSION

Access to reliable prescription drug information is essential to understanding potential drug interactions, side effects, and risks. Trusted online sources such as Medscape, WebMD, Drugs, and Drugbank were used for a thorough comparison of different drugs. This comprehensive analysis revealed specific interactions among specific drug pairs. It is worth mentioning that drug interactions between capecitabine, oxaliplatin and leucovorin have been identified in the studied websites.

### Oxaliplatin & <u>Capecitabine</u>

The risk or severity of adverse effects can be increased when Oxaliplatin is combined with Capecitabine. Immunosuppressive agents may exert an additive effect on other immunosuppressive agents, leading to a greater risk of infection due to bone marrow suppression.(160),(161)

### Leucovorin & Capecitabine

The serum concentration of Capecitabine can be increased when it is combined with Leucovorin. Leucovorin has been observed to increase the therapeutic effect of capecitabine, likely through its metabolite 5-fluorouracil (5-FU).5,1,2 Previous research have shown that leucovorin stabilizes the complex formed between the 5-FdUMP, a capecitabine metabolite, and thymidylate synthase, thus allowing 5-FdUMP to stay inside cancer cells for longer and exert its antineoplastic effect.3,4. Although the co-administration of leucovorin and capecitabine can increase the latter’s therapeutic effect due to prolonged exposure, it also increases the risk of capecitabine-associated side effects, such as severe enterocolitis, diarrhea, and dehydration.(162),(163),(164),(165)

## 5. CONCLUSION

In conclusion, rectal neoplasms present a complex challenge in oncology, necessitating tailored and effective treatment strategies. The study utilized the innovative RAIN protocol, integrating artificial intelligence to propose and evaluate drug combinations for rectal neoplasm therapy. Through systematic review and meta-analysis, the efficacy of Oxaliplatin, Leucovorin, and Capecitabine in treating rectal neoplasms was confirmed. These findings underscore the importance of ongoing research and implementation of advanced technologies to optimize patient care and treatment outcomes. Health policymakers are encouraged to prioritize the integration of AI-driven approaches like the RAIN protocol in healthcare policies, facilitating personalized and more effective management of rectal neoplasms for improved patient well-being.

### Abbreviations

STROBE: Strengthening the Reporting of Observational studies in Epidemiology
PRISMA: Preferred Reporting Items for Systematic Reviews and Meta-Analysis
RAIN: Systematic Review and Artificial Intelligence Network Meta-Analysis

## Authors’ contributions

### Nasrin Dashti

Data Gathering, Writing original draft. **Ali A. Kiaei:** Conceptualization, Methodology, Review & Editing, AI model implementation, Formal analysis, Supervision. **Mahnaz Boush:** Conceptualization, Methodology, AI model implementation. **Behnam Gholami-Borujeni:** Conceptualization, Investigation, Review & Editing.. **Alireza nazari:** Review & Editing, Validation, Review & Editing, Investigation.

## Funding

Not applicable.

## Availability of data and materials

Datasets are available through the corresponding author upon reasonable request.

## Ethics approval and consent to participate

Not applicable.

## Consent for publication

Not applicable.

## Conflict of interests

The authors have no conflict of interest.

## Notes

### Competing Interest Statement

The authors have declared no competing interest.

